# Axonal *T*_2_ estimation using the spherical variance of the strongly diffusion-weighted MRI signal

**DOI:** 10.1101/2021.08.19.456817

**Authors:** Marco Pizzolato, Mariam Andersson, Erick Jorge Canales-Rodríguez, Jean-Philippe Thiran, Tim B. Dyrby

## Abstract

In magnetic resonance imaging, the application of a strong diffusion weighting suppresses the signal contributions from the less diffusion-restricted constituents of the brain’s white matter, thus enabling the estimation of the transverse relaxation time *T*_2_ that arises from the more diffusion-restricted constituents such as the axons. However, the presence of cell nuclei and vacuoles can confound the estimation of the axonal *T*_2_, as diffusion within those structures is also restricted, causing the corresponding signal to survive the strong diffusion weighting. We devise an estimator of the axonal *T*_2_ based on the directional spherical variance of the strongly diffusion-weighted signal. The spherical variance *T*_2_ estimates are insensitive to the presence of isotropic contributions to the signal like those provided by cell nuclei and vacuoles. We show that with a strong diffusion weighting these estimates differ from those obtained using the directional spherical mean of the signal which contains both axonal and isotropically-restricted contributions. Our findings hint at the presence of an MRI-visible isotropically-restricted contribution to the signal in the white matter *ex vivo* fixed tissue (monkey) at 7T, and do not allow us to discard such a possibility also for *in vivo* human data collected with a clinical 3T system.

## 1. Introduction

One of the fundamental goals in magnetic resonance imaging (MRI) is the production of brain maps that provide, at each voxel, the quantification of the proportions and of the biophysical characteristics of the constituents of the brain’s tissue microstructure, such as neuronal cell bodies, axons, myelin, oligodendrocytes, astrocytes, etc.(Alexander, Dyrby, Nilsson and Zhang, 2019). This quantification holds the promise of identifying biomarkers that are sensitive to the presence and type of alterations and/or pathology. For this, it is fundamental to obtain specific measurements of the characteristics of each tissue constituent. Multi-compartmental biophysical modeling of the MRI signal has often been used for this purpose, however it entails challenges with regards to the estimation of the unknown values of the model parameters (Jelescu, Veraart, Fieremans and Novikov, 2016). To partially obviate this issue, research has focused on using particular acquisition regimes to measure a signal that retains the information from only a few key tissue constituents, such as the axons in the white matter tissue (Jensen, Glenn and Helpern, 2016; McKinnon and Jensen, 2019; Veraart, Nunes, Rudrapatna, Fieremans, Jones, Novikov and Shemesh, 2020). In particular, by using a strong diffusion weighting – generally summarized by a high b-value (*b*) – it is possible to measure a signal that almost entirely contains contributions from the compartments where water diffusion is more restricted. In this context, the term *restricted* is loosely used to indicate apparent low diffusivity and does not necessarily imply a specific type of diffusion process (White, McDonald, Farid, Kuperman, Karow, Schenker-Ahmed, Bartsch, Rakow-Penner, Holland, Shabaik et al., 2014). In white matter, since diffusion inside of axons is assumed to be more restricted than that outside of them (extra-axonal), a “sufficiently high” b-value leads to a diffusion signal for which the spherical mean over the acquired diffusion gradient directions mainly contains signal contributions from water spins inside of axons. In order to measure one key magnetic property of the axons, the axonal transverse relaxation time *T*_2a_, McKinnon and Jensen (2019) used this principle in combination with a transverse relaxation experiment. Their method consisted of observing the decay of the spherical mean of the strongly diffusion-weighted signal due to transverse relaxation, from which it is possible to estimate the axonal *T*_2_.

Microscopy studies have however revealed that axons may not be the only constituents of the white matter tissue microstructure that are characterized by a restricted diffusion process. For instance, Andersson, Kjer, Rafael-Patino, Pacureanu, Pakkenberg, Thiran, Ptito, Bech, Dahl, Dahl et al. (2020) illustrate the presence of vacuoles and of clusters of cell nuclei that have sizes compatible with restricted diffusion. Therefore, spins diffusing within these structures would still contribute to the strongly diffusion-weighted MRI signal, and would therefore bias the measurement of the axonal *T*_2_ calculated from the transverse relaxation of the spherical mean. Indeed, the presence of vacuoles and cell nuclei may be the source of a diffusion-restricted and isotropic contribution to the signal that survives at high b-values. It is therefore fundamental to have an estimator of the axonal *T*_2_ that is less biased by the presence of these isotropically-restricted signal contributions. This could enable a more accurate multi-compartmental modeling of the white matter tissue microstructure, where it is known that a small bias in the estimation of one parameter can cause biases in the estimation of all of the others. For instance, the assignment of an incorrect value of the axonal *T*_2_ can confound the estimation of anatomically relevant estimates such as the compartmental volume fractions, potentially having consequences for clinical interpretability. In addition, the comparison of the estimates from this unbiased estimator with those based on the spherical mean provides a new way to investigate the long-debated detectability of such isotropically-restricted signal contributions.

The presence of an isotropically-restricted signal contribution – possibly arising from spherical/cellular-like structures in the brain’s white matter – has been considered in the literature, particularly in the form of a “dot” signal contribution, corresponding to pools of apparently immobile water. In particular, the use of a dot compartment has been considered in biophysical modeling (Alexander, Hubbard, Hall, Moore, Ptito, Parker and Dyrby, 2010; Panagiotaki, Schneider, Siow, Hall, Lythgoe and Alexander, 2012; Ferizi, Schneider, Panagiotaki, Nedjati-Gilani, Zhang, Wheeler-Kingshott and Alexander, 2014; Zeng, Shi, Zhang, Ling, Dong and Jiang, 2018). However, the existence of this zero-diffusion contribution to the diffusion signal is debated: it has been deemed negligible in the context of clinical acquisitions (Dhital, Kellner, Kiselev and Reisert, 2018), or in non-fixed tissue (Veraart et al., 2020). Tax, Szczepankiewicz, Nilsson and Jones (2020) identified an isotropic and restricted (although less so than a dot) contribution to the strongly diffusion-weighted signal in some white matter regions.

Inspired by this debate, and with the aim of obtaining unbiased axonal *T*_2_ estimates, we propose to calculate the axonal transverse relaxation time from the directional spherical *variance* of the strongly diffusion-weighted signal which is, as we show, insensitive to the presence of isotropically-restricted contributions (dot or not). The *T*_2_ estimated from the spherical variance is indeed uniquely determined by anisotropic contributions to the measured directional signal, which in white matter are mainly represented by axons. Using data collected with a pre-clinical 7T MRI scanner from *ex vivo* fixed brain tissue of a Vervet monkey, and data collected with a clinical 3T MRI system from healthy human volunteers, we present evidence of the differences between the estimates of axonal *T*_2_ when using the spherical mean as compared to when unbiased estimates are obtained from the spherical variance.

## 2. Materials and methods

### 2.1. Spherical mean *T*_2_

The method proposed by McKinnon and Jensen (2019) uses a pulsed gradient spin echo (PGSE) (Stejskal and Tanner, 1965) sequence with b-value of 5000 or 6000 s/mm^2^ to collect data along different gradient directions and for two distinct echo times, *T E*_1_ and *T E*_2_. The indicated b-values are the results of a prediction based on the *in vivo* expected diffusivities at 37°C in the white matter and are chosen to ensure that the contribution of the extra-axonal water to the signal is negligible. More recently, a b-value of 4000 s/mm^2^ was deemed sufficient (Ramanna, Moss, McKinnon, Yacoub, Helpern and Jensen, 2020). In *ex vivo* scenarios, where the diffusivities are much lower, the b-value to choose has been indicated to be greater than or equal to 20000 s/mm^2^ (Veraart et al., 2020). Using such a high b-value, the axonal transverse relaxation was calculated by considering the spherical mean across the directions as

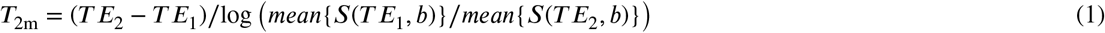

where the echo times need to be high enough to allow the contributions from myelin water to be neglected, and where

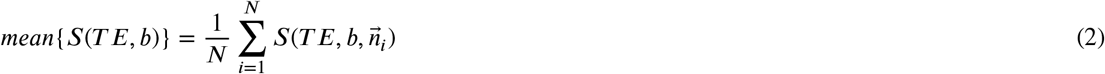

with 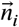 being the i-th gradient direction among the *N* acquired ones.

Equation 1 implicitly assumes that the only tissue constituents contributing to the strongly diffusion-weighted signal are axons. However, if in addition to axonal signal contributions there also are isotropically-restricted contributions coming, for instance, from cell nuclei and/or vacuoles, the estimated *T*_2_ would be biased.

### 2.2. Spherical variance *T*_2_

The spherical variance is the variance of signal samples acquired along many directions on a diffusion shell with fixed b-value. Intuitively, the variance is only influenced by contributions to the signal that vary directionally, i.e. it is not influenced by isotropic components as these only contribute to the mean of the signal. The directional signal measured in a voxel at a “high enough” b-value, e.g. *b* = 4000 mm^2^/s *in vivo* or *b* = 20000 mm^2^/s *ex vivo*, can be formulated as

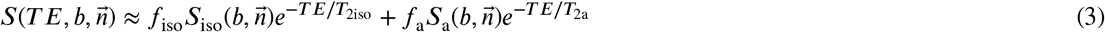

where for simplicity only one isotropic (iso) and one axonal (a) compartment are considered with the respective signal fractions (volume fractions modulated by proton density and longitudinal relaxation), *f*_iso_ and *f*_a_, diffusion attenuations, *S*_iso_(·) and *S*_*a*_ (·), and transverse relaxation times, *T*_2iso_ and *T*_2a_. The sample spherical variance is then formulated as

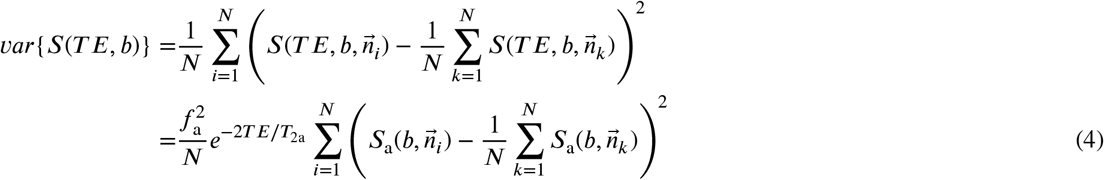

where we have used the fact that an isotropic compartment displays a signal decay in each direction that is equal to the average across all directions, that is

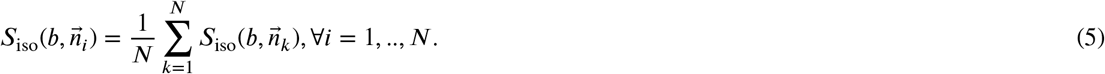

The relation in eq. 4 leads to the estimation of the axonal transverse relaxation time without the influence of isotropic compartments as

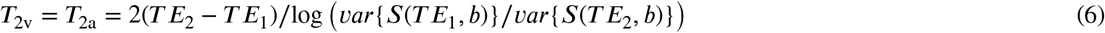

where all the unknowns in eq. 4 cancel each other when dividing the spherical variances for two different echo times (at fixed b-value), except for the exponential terms that account for the dependency on the axonal *T*_2a_.

When assuming the presence of isotropic compartments that contribute to the spherical mean signal, eq. 1 is no longer a valid estimator of the axonal transverse relaxation time, *T*_2a_, as it assumes an “axon-only” model of the strongly diffusion-weighted signal. The use of eq. 6, based on the spherical variance, leads to an unbiased estimation of the axonal *T*_2_ also when isotropic compartments are present because it only accounts for anisotropic contributions to the strongly diffusion-weighted signal, such as those coming from axons.

The variance of the signal is however more sensitive to noise. The noise variance, in the case of zero-mean Gaussian distributed noise with variance *σ*^2^, is an additive term to the true variance, i.e the measured variance is *var*{*S*(*T E, b*)} + *σ*^2^. In the absence of a perfect knowledge of the noise variance, it is therefore essential to perform denoising to attenuate noise before using eq. 1. To mitigate the issues related to the additive noise variance, should the denoising not remove the noise entirely, it is convenient to use approximation and eventually regularization. A typical strategy for doing so is to approximate the signal on a shell with spherical harmonics, and eventually apply Laplace-Beltrami regularization (Descoteaux, Angelino, Fitzgibbons and Deriche, 2007). The spherical harmonics expansion of the signal is

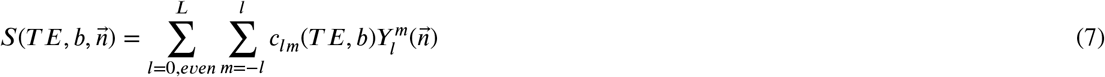

where 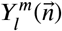 are the real spherical harmonics basis functions of order *l* and *m* (see appendix A) – the *l*-order subscript only admits even values, corresponding to even spherical harmonics, because of the antipodal symmetry of the diffusion signal – and where *c*_*lm*_(*T E, b*) are the corresponding coefficients. Laplace-Beltrami regularization is particularly suitable for this kind of applications as the higher order coefficients, associated to high frequency oscillations such as noise, are more penalized than lower order coefficients. The use of regularization implies selecting the amount of smoothing, which is typically expressed by a meta-parameter, *λ*, (Descoteaux et al., 2007). The spherical variance used to evaluate eq. 6 can then be calculated as (Zucchelli, Deslauriers-Gauthier and Deriche, 2020)

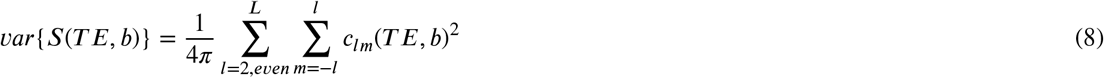

which only uses the coefficients with *l* ≥ 2 thus excluding the zeroth coefficient that only encodes for the spherical mean of the signal.In addition to noise attenuation, the use of the spherical harmonics expansion offers the advantage of allowing more flexible acquisitions with regards to the diffusion gradient directions.

Looking at eq. 3 it appears evident that to estimate the axonal transverse relaxation time, *T*_2a_, one could remove the spherical mean from the overall signal, to remain with only a reminder of the leftover anisotropic signal contributions. With that, it would be possible to perform a *T*_2_ estimation for each pair of signal samples collected with the same b-value along collinear directions at different echo times. This clearly constrains the acquisitions to be performed along the same directions, and would directly suffer from any misregistration errors across the collected images. Nevertheless, this approach leads to more instability or more bias in the presence of residual noise after denoising compared to using the spherical variance (see appendix B).

### 2.3. Interpreting the *T*_2_ estimates

The b-value used for the acquisition and the type of estimation (e.g. mean or variance) have a direct impact on which constituents of the tissue microstructure contribute to the *T*_2_ estimate. Equation 1 can be used to calculate the *T*_2_ at *b* = 0, which is referred to as the *b* = 0 average *T*_2_, or at “high” b-value, i.e. *T*_2m_. Equation 6 is used to estimate the *T*_2_ only at a non-zero b-value, *T*_2v_. In white matter, these different estimates of the transverse relaxation time differ in that

- the spherical variance *T*_2v_ calculated at *high* b-value accounts only for anisotropic, e.g. axonal, contributions from restricted compartments;
- the spherical variance *T*_2v_ calculated at *lower* (non-zero) b-value additionally accounts for the anisotropic contributions from less restricted anisotropic compartments, e.g. the extra-axonal water;
- the spherical mean *T*_2m_ calculated at *high* b-value accounts for all contributions from restricted compartments, e.g. from both axons and cell nuclei and/or vacuoles (or generally isotropically-restricted compartments);
- the spherical mean *T*_2m_ calculated at *lower* (non-zero) b-value additionally accounts for contributions from isotropic compartments where diffusion is not (or is less) restricted, and from the extra-axonal water;
- the *b* = 0 average *T*_2_ accounts for similar contributions to those determining the spherical mean *T*_2m_ calculated at lower b-value with the addition of even less restricted compartments.

We finally note that the *T*_2_ estimates based on the spherical mean, unlike those based on the spherical variance, can be more affected by partial volume contamination with isotropic signal contributions from the gray matter (GM) and the cerebrospinal fluid (CSF).

### 2.4. Ex vivo data

Data was collected from an *ex vivo* fixed Vervet (Chlorocebus aethiops) monkey brain, obtained from the Montreal Monkey Brain Bank. The monkey, cared for on the island of St. Kitts, had been treated in line with a protocol approved by The Caribbean Primate Center of St. Kitts. The brain had previously been stored and prepared according to Dyrby, Baaré, Alexander, Jelsing, Garde and Søgaard (2011). The data was collected with a Bruker Biospec 70/20 7.0 T scanner (Billerica, Massachusetts, USA) using a quadrature RF coil (300MHz). The brain was let to reach room temperature and to mechanically stabilize prior the start of the acquisition. The acquisition was conducted using a constant air flow directed towards the brain to avoid short-term instability artifacts (Dyrby et al., 2011). A single-line readout PGSE sequence with pulse duration *δ* = 9.6 ms and separation Δ = 17.5 ms was used to collect the data organized in five shells with *b* = 4000 s/mm^2^, 7000 s/mm^2^, 23000 s/mm^2^, 27000 s/mm^2^, and 31000 s/mm^2^ each containing the same 96 directions which were obtained by electrostatic repulsion (Jones, Horsfield and Simmons, 1999), plus *b* = 0 images. One dataset was collected at echo time *T E* = 35.5 ms and the other at *T E* = 45.5 ms. We expect mild contamination from myelin at these echo times. Assuming a myelin *T*_2mye_ = 11 ms (Birkl, Doucette, Fan, Hernández-Torres and Rauscher, 2021) and an axonal *T*_2a_ = 30 ms, at *T E* = 35.5 ms a ratio between myelin volume fraction and axonal volume fraction of 1/2 (g-ratio ≈ 0.82) would lead to approximately a 6% myelin signal contribution, while a ratio of 1/4 (g-ratio ≈ 0.89) would lead to an approximately 3% myelin signal contribution. At *T E* = 45.5 ms the expected myelin signal contributions reduce to approximately 3.5% and 1.8% respectively. The percentage contributions of the myelin water are actually expected to be lower than those values since the myelin transverse relaxation time reported previously is related to *in vivo* tissue at 3T and it might decrease with increased field strength (7T) and due to the lower temperature and to the formalin fixation (Birkl, Langkammer, Golob-Schwarzl, Leoni, Haybaeck, Goessler, Fazekas and Ropele, 2016). The estimated SNR in WM was around 45. Images were collected with a repetition time of *T R* = 3500 ms, at a 0.5 mm isotropic resolution, and with an image matrix of 256×128×30 voxels. Total scan time was of approximately 6 days.

### 2.5. Human clinical data

Data was collected from a 27 year old female (sbj1) and a 34 year old male (subj2) healthy volunteer. For the latter, a retest was performed (sbj2r) two weeks after the first acquisition. The acquisition and the protocol were authorized by Danish ethical commission of Region Hovedstaden (Journal-nr.: H-21022514). Data was acquired using a Siemens 3T Prisma MRI system (Siemens Heathineers, Erlangen, Germany) using a 64-channel head coil. A monopolar pulse (PGSE) was used for collecting data organized in two shells with *b* = 1000 s/mm^2^ and 5000 s/mm^2^, each containing the same 96 directions which were obtained as done for the *ex vivo* data. Non-weighted (*b* = 0) images were collected with reversed phase-encode blips, resulting in pairs of images with distortions going in opposite directions for the correction of the susceptibility-induced off-resonance field in the whole dataset. One dataset was collected at echo time *T E* = 80 ms and the other at *T E* = 89 ms. For the shell with *b* = 5000 s/mm^2^ an additional dataset was acquired with *T E* = 85 ms. Finally, another shell with *b* = 2500 s/mm^2^ and *T E* = 80 ms was acquired for the estimation of the fiber orientation distribution function (fODF) to be used in the analysis. The MRI scanner automatically adapts the pulse length and separation of the PGSE sequence depending on the prescribed echo time as also reported in McKinnon and Jensen (2019). Therefore, the echo time range [80,89] ms was chosen as it minimizes the difference in diffusion time between the *b* = 5000 s/mm^2^ data collected with different echo times while still guaranteeing an observable decay due to relaxation. Images were collected with a repetition time of *T R* = 3840 ms, at a 2.3 mm isotropic resolution, multi-band factor 3, and with an image matrix of 88×90×74 voxels. The total scan time was of approximately 56 minutes per subject.

### 2.6. Data processing

The data was denoised using the method described by Ma, Uğurbil and Wu (2020) with a window size of 3 voxels. This denoising is based on the estimation of the noise variance in association with a Rician variance stabilization technique (Foi, 2011), after which optimal shrinkage with respect to the mean squared error (Gavish and Donoho, 2017) is applied to the singular values extracted from local 3D isotropic patches of data with side determined by the chosen window size. The denoising removes the Rician bias and increases the SNR of the images. After denoising, Gibbs ringing removal according to Kellner, Dhital, Kiselev and Reisert (2016) was applied using the implementation available in MRtrix3^1^. While for the *ex vivo* data a visual inspection revealed that image registration was not necessary, for the *in vivo* human data a registration was performed using FSL’s eddy (Andersson and Sotiropoulos, 2016) after using FSL’s topup which was used to estimate the susceptibility-induced off-resonance field from the pairs of *b* = 0 images collected with reversed phase-encode blips (Andersson, Skare and Ashburner, 2003; Smith, Jenkinson, Woolrich, Beckmann, Behrens, Johansen-Berg, Bannister, De Luca, Drobnjak, Flitney et al., 2004).

The evaluation of eqs. 1, 2, 4, and 6 was carried out using NumPy^2^ (Harris, Millman, van der Walt, Gommers, Virtanen, Cournapeau, Wieser, Taylor, Berg, Smith, Kern, Picus, Hoyer, van Kerkwijk, Brett, Haldane, Fernández del Río, Wiebe, Peterson, Gérard-Marchant, Sheppard, Reddy, Weckesser, Abbasi, Gohlke and Oliphant, 2020). When multiple echo times were available, the estimation of the *T*_2_ values was performed by using the SciPy^3^ (Virtanen, Gommers, Oliphant, Haberland, Reddy, Cournapeau, Burovski, Peterson, Weckesser, Bright, van der Walt, Brett, Wilson, Millman, Mayorov, Nelson, Jones, Kern, Larson, Carey, Polat, Feng, Moore, VanderPlas, Laxalde, Perktold, Cimrman, Henriksen, Quintero, Harris, Archibald, Ribeiro, Pedregosa, van Mulbregt and SciPy 1.0 Contributors, 2020) implementation of differential evolution (Storn and Price, 1997) to minimize objective functions corresponding to eqs. 1 and 6 while considering the mean squared error over the possible non-repeated combinations of echo times for *T E*_1_ and *T E*_2_. The visualization of the results was performed with Matplotlib^4^ (Hunter, 2007).

### 2.7. Synthetic data

Synthetic data was generated based on the acquisition protocol for the *ex vivo* acquisition, with diffusion parameters set to values in line with fixed tissue at room temperature. To simulate the data we first estimated the tissue signal fractions for white matter (WM), gray matter (GM), and cerebrospinal fluid (CSF) using the multi-shell multi-tissue constrained spherical deconvolution framework (Jeurissen, Tournier, Dhollander, Connelly and Sijbers, 2014) in its generalized version implemented in the Dmipy^5^ library (Fick, Wassermann and Deriche, 2019). The fiber orientation distribution function (fODF) associated to the WM compartment was estimated with spherical harmonics up to order *L* = 8, using the shells with *b* = 4000 s/mm^2^ and *b* = 7000 s/mm^2^. The signal fractions were used to regenerate the synthetic brain dataset. For WM, three compartments were simulated: an axonal compartment with volume fraction of 0.6, an extra-axonal compartment with volume fraction 0.3, and an isotropically-restricted (spherical) compartment with volume fraction of 0.1. The axonal and extra-axonal compartments were simulated with axisymmetric tensors. The axonal compartment had parallel and perpendicular diffusivities of 2*e* − 10*mm*^2^/*s* and 3.5*e* − 11*mm*^2^/*s*, and the extra-axonal one of 6*e* − 10*mm*^2^/*s* and 1.5*e* − 10*mm*^2^/*s* respectively. Values were chosen to ensure a more rapid decay of the extra-axonal spherical mean signal as compared to the axonal one. Because of the unknown nature of some parameter values, some informed guesses have guided our choices. However, the exact values of the simulated parameters do not significantly affect the subsequent analysis which can be considered general with respect to the axonal and isotropic transverse relaxation times. The isotropically-restricted compartment was simulated as isotropic Gaussian diffusion with diffusivity 1*e*−11*mm*^2^/*s*. GM and CSF were simulated as isotropic Gaussian diffusion components with diffusivities 7*e*−10*mm*^2^/*s* and 9*e*−10*mm*^2^/*s* respectively. The transverse relaxation times were fixed to 30 ms (axonal), 50 ms (extra-axonal), 80 ms (GM), and 500 ms (CSF). The isotropically-restricted compartment was simulated with *T*_2iso_ = 50 ms. Rician noise was applied to the ground-truth data according to a signal-to-noise ratio (SNR). This was defined as the ratio between the median value of the intensity of the *b* = 0 volume over the white matter and the standard deviation of the additive complex Gaussian noise from which the Rician volumes are generated. Two SNRs were tested, SNR=40 and SNR=20. The synthetic datasets were then denoised using the same strategy used for the acquired data and no further processing was performed.

## 3. Results

### 3.1. Simulations

The experiments based on synthetic data are designed to illustrate the potential bias arising from estimating of the axonal transverse relaxation time *T*_2a_ based on the spherical mean (eq. 1) in the presence of an isotropically-restricted compartment, and to evaluate the robustness of the estimates based on the spherical variance. Fig. 1 illustrates the obtained *T*_2_ maps (*T*_2a_ = 30 ms) calculated at *b* = 23000 s/mm^2^. The changes across the maps obtained for different SNRs highlight the impact of the residual noise (after denoising) which leads to a high variability in the estimates.

**Figure 1:**
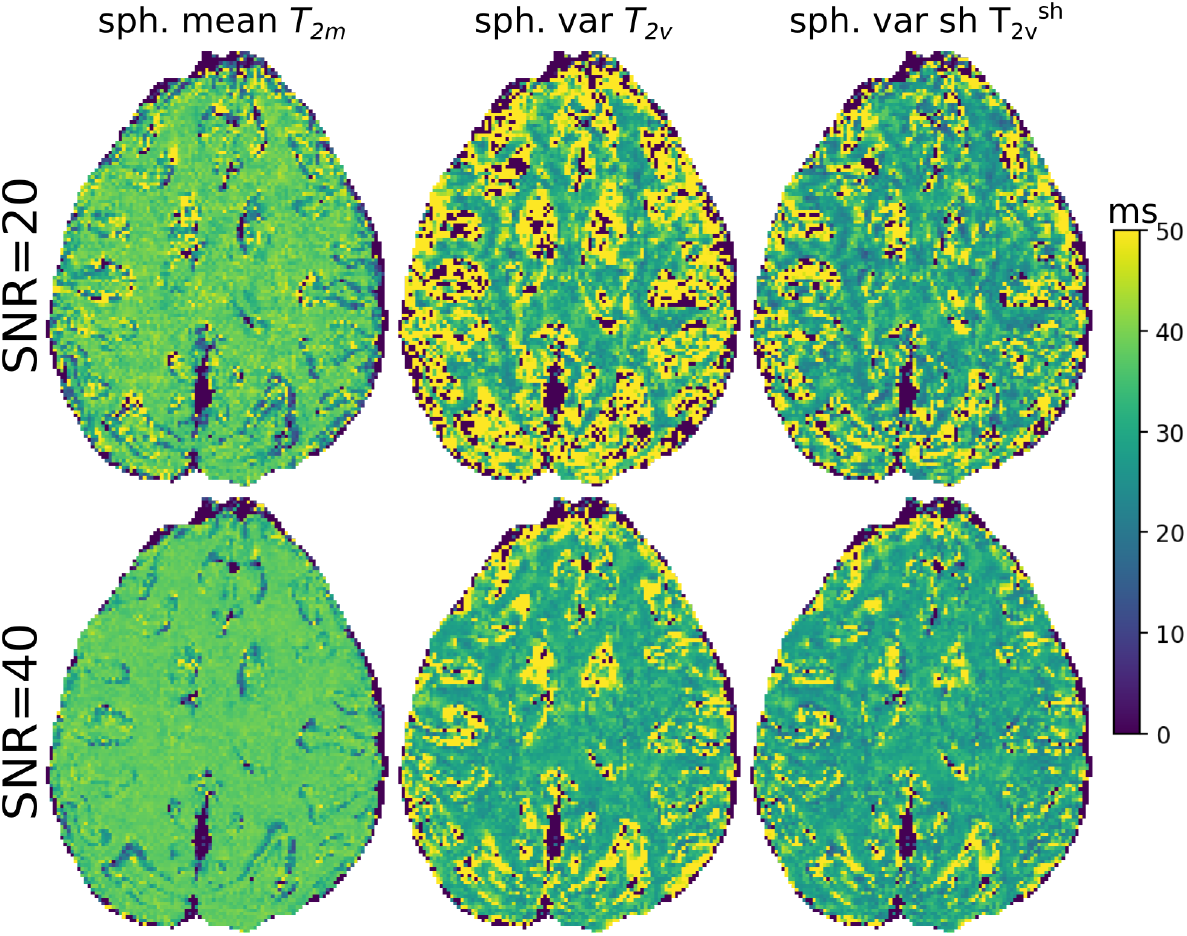
Synthetic data. Maps of axonal *T*_2_ estimates (ms) calculated with different methods at *b* = 23000 s/mm^2^ for SNR 20 and 40 on simulated data (section 2.7). The left-most column reports the *T*_2_ estimates based on the spherical mean, the central one those based on the spherical variance, and the right-most one the estimates based on the spherical variance estimated using the spherical harmonics expansion (*L* = 10) with a Laplace-Beltrami regularization amount of *λ* = 0.003. The ground-truth axonal *T*_2a_ is 30 ms.

The histograms shown in fig. 2 reveal that the spherical mean (*T*_2m_) estimates are centered around a value in between the axonal transverse relaxation time and that of the isotropically-restricted compartment. This indicates that the bias is induced by the presence of the isotropically-restricted compartment which, despite the small volume fraction, has a relevant signal contribution at high b-value. Indeed, the spherical variance estimates (*T*_2v_) are centered closer to the axonal *T*_2a_ value. On the other hand, the spherical mean *T*_2_ estimates show a smaller variance compared to the spherical variance *T*_2_ estimates, indicating a higher stability.

**Figure 2:**
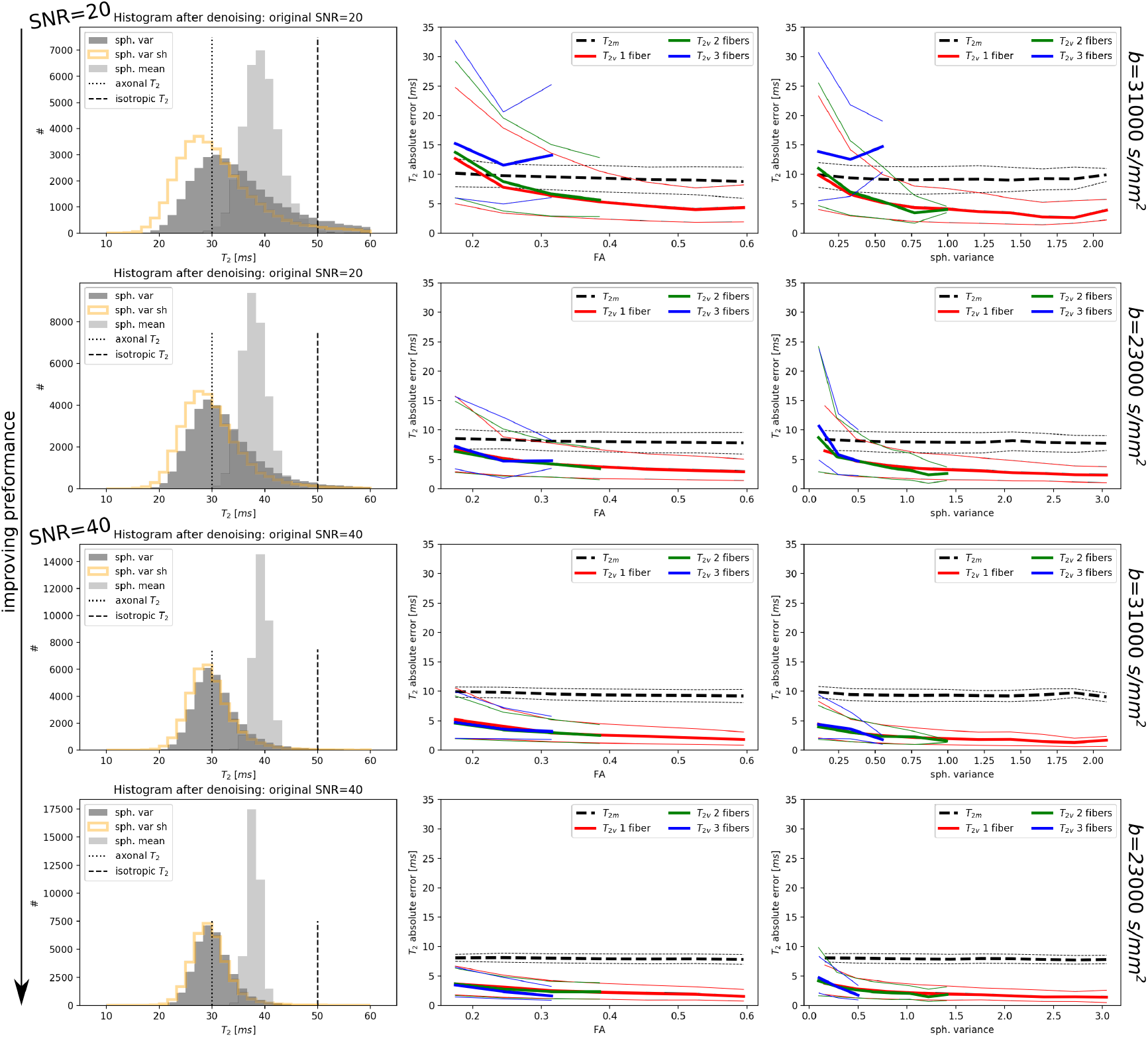
Performance on synthetic data for various estimators of the axonal transverse relaxation time for SNR 20 and 40 and for different b-values (*b* = 31000 s/mm^2^ and *b* = 23000 s/mm^2^). The histograms compare the distributions of the estimates for the spherical variance (eventually regularized using Laplace-Beltrami with *λ* = 0.003 and spherical mean estimators. The values of the axonal and isotropic compartment transverse relaxation times are indicated with vertical lines. Plots on the right hand side illustrate the absolute error in the axonal *T*_2_ estimation achieved with the spherical variance (non-regularized), *T*_2v_, and with the spherical mean, *T*_2m_. Thick lines represent the median error, whereas the thinner lines indicate the 25th and 75th percentiles of the absolute error distributions. In the case of the spherical variance, the error lines are color coded depending on the number of fibers detected in the voxels. The error is plotted as a function of the ground-truth values of the FA (second column) and of the spherical variance (third column) of the synthetic data. The performance of all estimators increase as the SNR increases and as the b-value reduces.

The accuracy (mode of the distribution) and stability (variance of the distribution) of the spherical variance estimator improve as the SNR increases and as the b-value decreases (from 31000 s/mm^2^ to 23000 s/mm^2^ as shown in fig. 2). The improved performance as the b-value reduces is explained by an increased spherical variance of the signal as compared to the variance contribution arising from the residual noise. A weaker diffusion weighting incurs less signal decay, thus the noise variance – which is substantially constant across b-values – represents a smaller fraction of the total variance of the signal and therefore has less impact. This can be observed from the plots of the absolute *T*_2_ estimation error on the right side of fig. 2. The absolute error reduces with the increasing ground-truth variance of the signal (third column) and this in turn increases when reducing the b-value (see differences in the x axis coordinate values between the cases of *b* = 31000 s/mm^2^ and *b* = 23000 s/mm^2^). As shown in the second column of fig. 2, similar results are obtained while observing the trend of the error with respect to fractional anisotropy (FA) (Basser, Mattiello and LeBihan, 1994) – correlated to the spherical variance (Zucchelli et al., 2020) – which was calculated using the lower shells (*b* = 4000 s/mm^2^ and *b* = 7000 s/mm^2^) of the ground-truth data with Dipy ^6^ (Garyfallidis, Brett, Amirbekian, Rokem, Van Der Walt, Descoteaux and Nimmo-Smith, 2014).

Since the spherical variance of the ground-truth signal is expected to considerably reduce in the presence of multiple fibers with different orientations, we classified the white matter voxels based on the number of detected fibers using the constrained spherical deconvolution method (Tournier, Calamante and Connelly, 2007) and peak detection implemented in Dipy using a minimum peaks separation angle of 25° as typically done. As expected, the spherical variance is lower for voxels with multiple fibers and, consequently, the error is higher. It is important to note that this is purely an effect due to the presence of residual noise. In other words, as long as the spherical variance is different from zero it is possible to accurately measure the axonal *T*_2_ on noise-free data.

The use of Laplace-Beltrami regularization for computing the spherical variance (first column of fig. 2) is beneficial for the stability but can result in a mild bias as seen from the modes of the distributions that are slightly misaligned with *T*_2*a*_. It is non-trivial to choose an amount of regularization in this setup. We found that the use of spherical harmonics, without regularizing, suffers less from this biasing effect. Moreover, depending on the selected order of the expansion, the use of spherical harmonics can provide an approximation of the directional signal which can diminish the effects of the residual noise. In the light of these considerations, this last strategy will therefore be adopted for the *in vivo* data.

Overall, the simulation results suggest that to estimate the axonal transverse relaxation time, a trade-off between bias and variance of the estimators is necessary. While the spherical variance *T*_2_ estimator is less biased in the presence of isotropically-restricted compartments, the corresponding estimates suffers from a higher variability. Moreover, higher reliability of the estimates is achieved in voxels where the noise-free spherical variance is higher, i.e. where the presence of residual noise has less impact. Therefore, estimates are more reliable in voxels with single fiber bundles, low orientational dispersion, and at lower b-values.

### 3.2. Ex vivo data

The trends of the *T*_2_ estimates are analyzed as a function of the values of other diffusion descriptors derived from DTI. In particular, we focus on fractional anisotropy (FA) and on the angle between the main diffusion orientation and the B0 field direction, similarly to what previously done by McKinnon and Jensen (2019). The value of FA is influenced by several factors, such as the presence of multiple fibers, the presence of cell-like/spherical isotropic compartments, and the partial voluming with isotropic tissue components such as gray matter (GM) and cerebrospinal fluid (CSF). In the analysis we try to isolate only the dependency of FA on the presence of cell-like/spherical isotropic compartments by a) performing the analysis on a conservative white matter mask to minimize the probability of partial voluming effects; b) using high b-value data when possible, such that GM and CSF contributions attenuate almost entirely; c) including voxels where only a single fiber is detected.

#### 3.2.1. Average, spherical mean, and spherical variance T_2_

The second and third columns of fig. 3 illustrate the maps of the axonal T2 obtained, at high b-value, from the spherical mean (*T*_2m_) and from the spherical variance (*T*_2v_), along with their differences (last two columns). The figure also illustrates the average *T*_2_ calculated on the *b* = 0 data (first column) using eq. 1. The *T*_2v_ maps unravel more details compared to both the average *T*_2_ and to *T*_2m_. In these maps, indeed, partial volume effects often “hide” the axonal *T*_2_ value. The difference maps also reveal that the order of the *T*_2v_ and *T*_2m_ estimates changes regionally.

**Figure 3:**
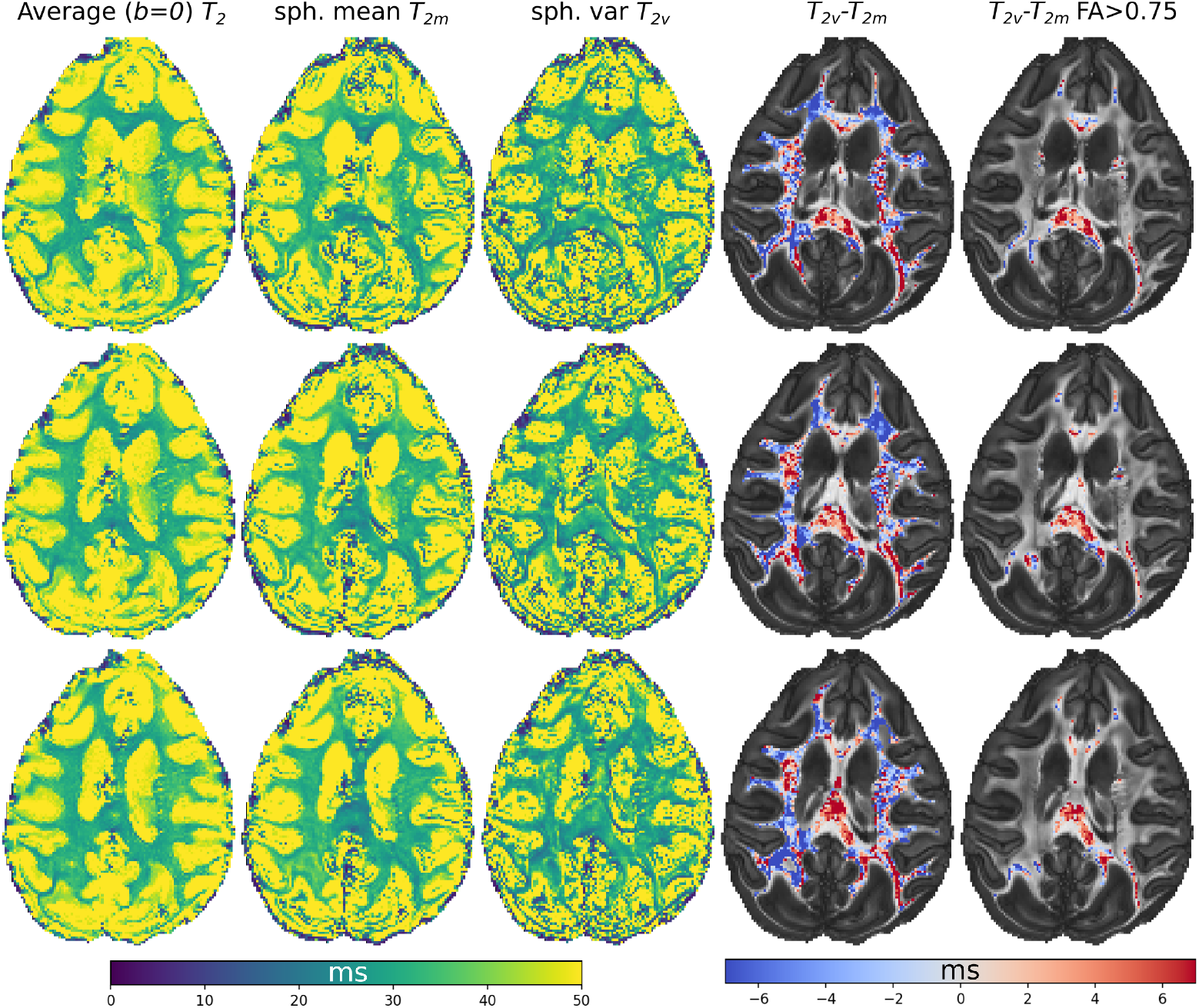
*Ex vivo* data. The average (*b* = 0), spherical mean (*T*_2m_, *b* = 23000 s/mm^2^) and spherical variance (*T*_2v_, *b* = 23000 s/mm^2^) transverse relaxation time estimates (ms), and the difference maps (last two columns) in a conservative white matter mask, eventually considering only voxels with FA > 0.75 (last column). Three different axial slices are shown (rows).

#### 3.2.2. Regional dependency of T_2_ estimates

In regions of high fractional anisotropy (e.g. FA > 0.75) the spherical variance leads to higher *T*_2_ estimates. From fig. 4b, c and d it is possible to observe that as FA increases, the spherical mean *T*_2_ estimates shift towards the values of the spherical variance estimates and slightly underestimate them for FA ≈ 0.75. Globally, however, the spherical variance estimates are slightly lower than those from the spherical mean, as illustrated in fig. 4a which reports histograms calculated over a conservative white matter mask. The variance of the distribution is larger for the spherical variance estimates, possibly because of the presence of residual noise. Supplemental fig. S1 illustrates a more complete picture where the histogram of the spherical variance, mean, and average *T*_2_ estimates are reported for each FA range and number of detected fibers. While observing that at low FA values the spherical variance estimates seem to be more unstable, at FA ≈ 0.2 the trends described for the single fiber voxels in the bottom histograms of fig. 4 are also verified for the voxels with multiple fibers, (third and fourth columns of fig. S1), and globally for the whole white matter (first column of fig. S1). These results summarize what illustrated in the last column of fig. 3

**Figure 4:**
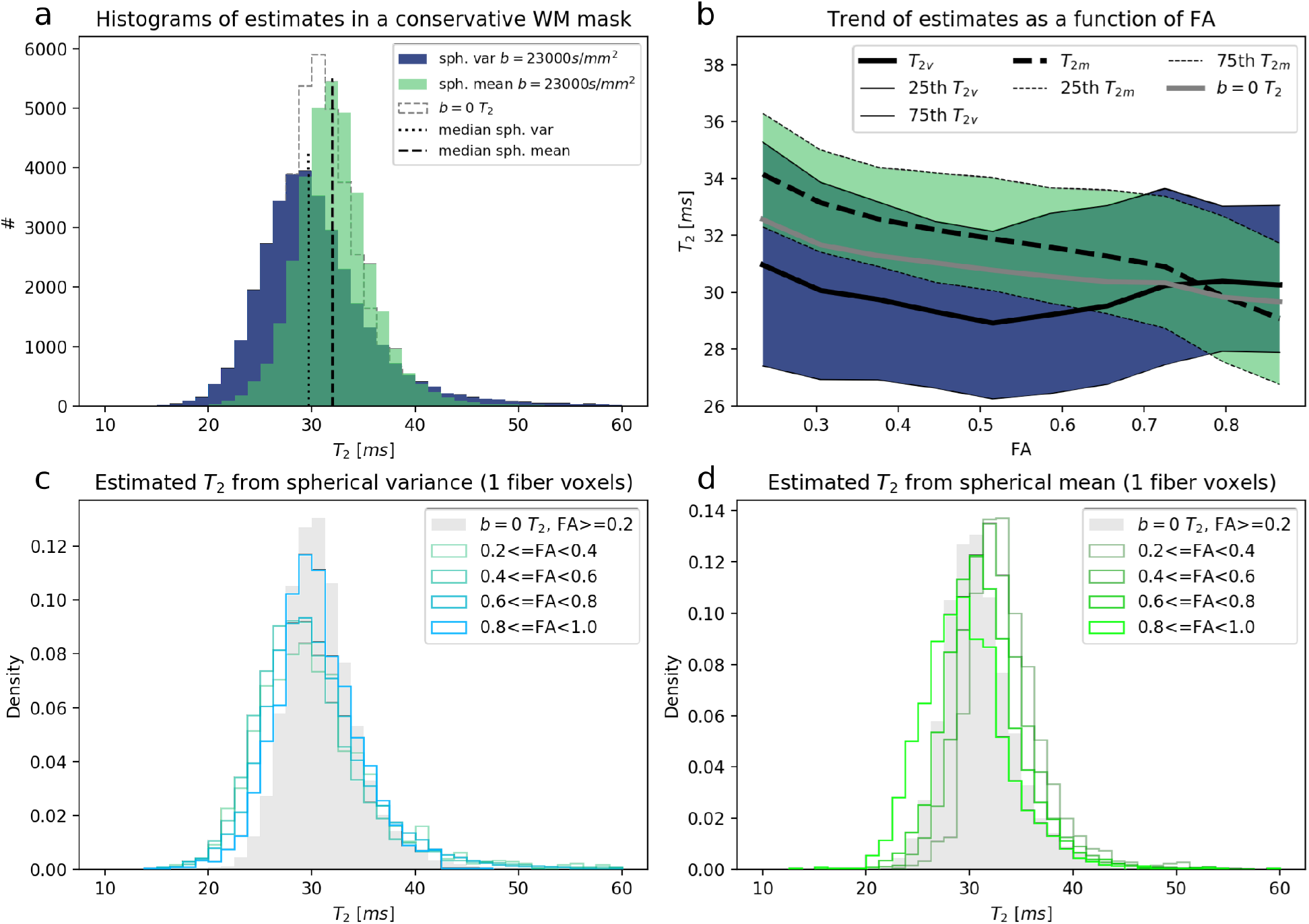
*Ex vivo* data. Histograms of the average (*b* = 0), spherical variance (*T*_2v_) and spherical mean (*T*_2m_) transverse relaxation time calculated within a conservative white matter mask. The spherical variance and spherical mean estimates were calculated at *b* = 23000 s/mm^2^ (top left). Variations of estimates as a function of FA (top right). Bottom, histograms as in the top left image calculated for different FA thresholds for the spherical variance (left) and spherical mean (right).

The results illustrated in fig. 4b show that the spherical mean *T*_2m_ estimates have a clear decreasing trend as a function of FA, while such a trend is less clear for the spherical variance estimates, *T*_2v_. The change of *T*_2m_ estimates as a function of FA could be induced by two factors: a) the values of FA in white matter voxels with only one fiber are determined by the change in the quantity of isotropic compartments or by the change in axonal orientational dispersion, either of which is also causing the change in the measured *T*_2_; b) the values of higher FA are associated with regions of the brain where the fibers present a specific alignment with respect to the B0 field direction (e.g. 90° in the corpus callosum) thus causing a lower value of the estimated *T*_2_ due to susceptibility artifacts, which can be particularly relevant at 7T (Sati, Silva, van Gelderen, Gaitan, Wohler, Jacobson, Duyn and Reich, 2012): although this should influence equally spherical mean and variance based estimates, there might be some unexpected differences. In fig. 5 the median values of the estimated *T*_2_ distributions from both the spherical mean and variance are plotted as a function of the detected angle between the main diffusion direction (calculated with DTI) and the direction of the B0 field, and for different ranges of FA values (top-right image). For completeness, the histograms of the direction as a function of FA is reported in the top-left corner of fig. 5. The localization of the selected voxels based on the selected FA ranges is shown in the maps at the bottom of the same figure. Of these voxels, only those where only one fiber was detected were used. This was done to exclude that the trend could be caused by substantial differences of axonal *T*_2_ for different bundles crossing within voxels, since the presence of such differences was suggested before (Barakovic, Tax, Rudrapatna, Chamberland, Rafael-Patino, Granziera, Thiran, Daducci, Canales-Rodríguez and Jones, 2021).

**Figure 5:**
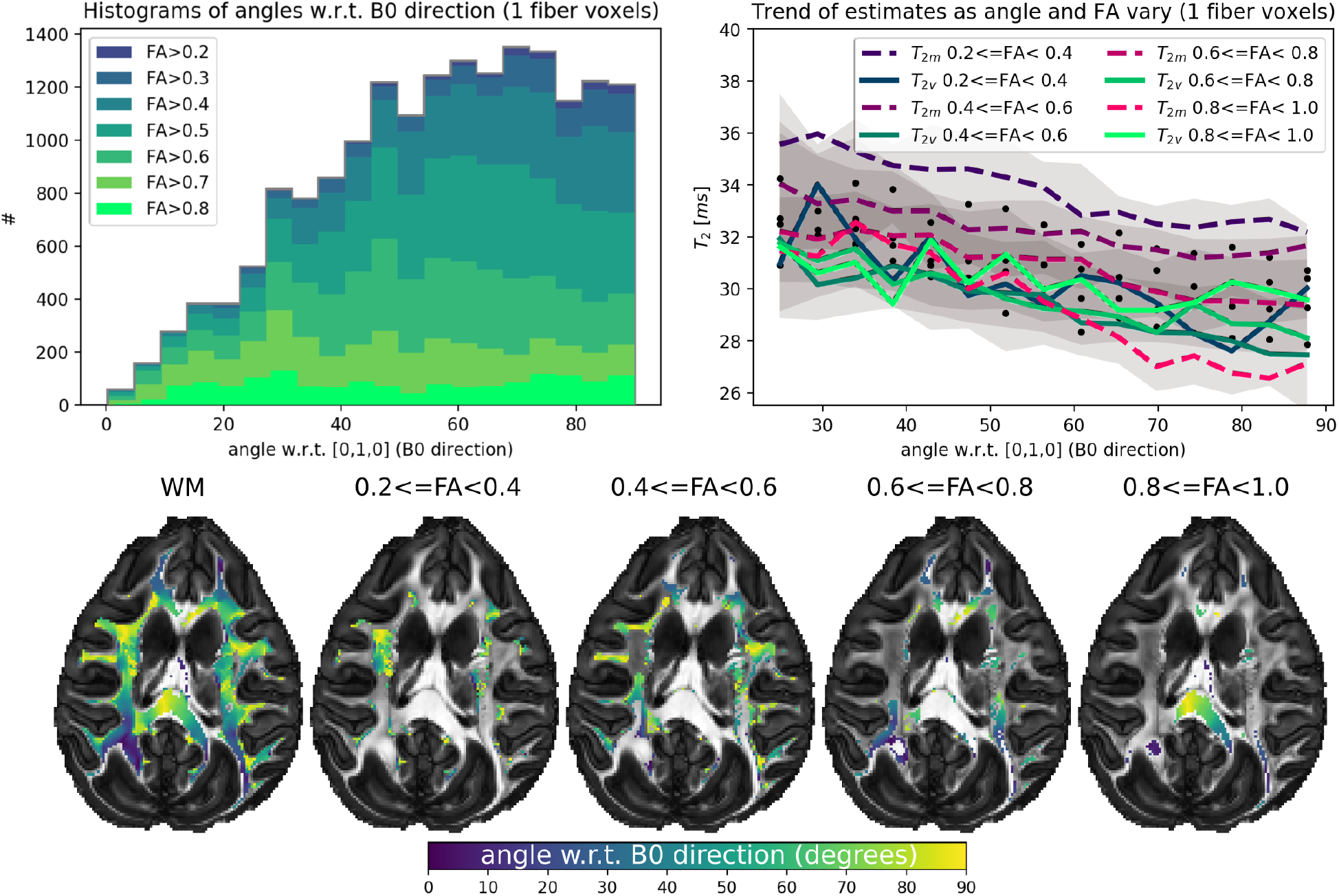
*Ex vivo* data. In the top-right, the dependency of the spherical mean (*T*_2m_, *b* = 23000 s/mm^2^), spherical variance (*T*_2v_, *b* = 23000 s/mm^2^), and average (*b* = 0) *T*_2_ estimates (dots) as a function of the angle between the main diffusion direction (estimated with DTI) and the direction of the B0 field, shown for different ranges fractional anisotropy (FA) values. Dashed lines correspond to the median values for *T*_2m_, continuous lines to *T*_2v_, whereas the dots indicate the median values of the *b* = 0 average estimates. The gray shaded areas indicate the 25th to 75th percentile ranges corresponding to the latter estimates. On the left, the histograms of the angles for different ranges of FA. In the bottom, maps show the detected angle (in degrees) overlaid to the FA map in WM (left-most map) and for different ranges of FA.

The results confirm the presence of a dependency of *T*_2_ values with respect to the angle between the fibers and the main magnetic field direction. Indeed, the median values of the estimates decrease of about 2 − 3 ms while passing from a 30° to a 90° angle. Nevertheless, the spherical mean *T*_2m_ estimates appear to have a stronger dependency on FA than on fiber orientation, with the estimated median *T*_2_ for low/high FA values differing by as much as 6 ms. This does not seem to be the case (or at least it is to a lesser extent) for the spherical variance *T*_2v_ estimates. Therefore, changes in *T*_2m_ appear to be related to changes in FA, which in turn could be determined by the change in quantity of isotropic compartments or by differences in the axonal orientational dispersion. However, the spherical variance *T*_2v_ estimates, which are insensitive to the presence of isotropic compartments but equally affected by orientational dispersion, show less variability with respect to FA. Hence, the difference in the extent of the variations of *T*_2m_ and *T*_2v_ estimates with respect to FA could better be explained by the presence of isotropic compartments rather than the occurrence of orientational dispersion of axons.

#### 3.2.3. The extra-axonal compartment

The *T*_2_ of the extra-axonal water may be qualitatively observed by relating the average (*b* = 0) *T*_2_ estimates with those obtained via the spherical mean and the spherical variance estimators at *b* = 23000 s/mm^2^. In white matter, the *b* = 0 *T*_2_ is determined by all of the compartments (axonal, isotropic, and extra-axonal). In regions characterized by a medium range FA (0.2 ≤ FA < 0.8), the average *b* = 0 *T*_2_ estimates have values that are intermediate between the lower spherical variance and the higher spherical mean ones (fig. 4b). Since the high b-value spherical mean and the average (*b* = 0) *T*_2_ estimates should mainly differ by the contributions of the extra-axonal water, the results suggest that the extra-axonal water has lower transverse relaxation times than the combination of axonal and isotropically-restricted compartments.

A more direct way to determine the order of *T*_2_ values between the axonal and the extra-axonal water is to observe the spherical variance estimates as a function of the b-value. Indeed, the estimates of the spherical variance *T*_2_ at a low b-value should be contaminated by anisotropic contributions from the extra-axonal water. In other words, the spherical variance *T*_2_ estimates calculated at *b* = 4000 s/mm^2^ will account for both the axonal and extra-axonal water contributions (both anisotropic), while the estimates calculated at *b* = 23000 s/mm^2^ should only account for the axonal water contribution. The right-side histograms in fig. 6 illustrate the distributions of the spherical variance *T*_2v_ estimates as a function of the b-value. These were also calculated only accounting for WM voxels where only one fiber was detected. While illustrating that there substantially are no differences between the estimates in the high b-value regime (for *b* = 23000, 27000, and 31000 s/mm^2^), the results show that at *b* = 4000 s/mm^2^ the estimated variance-based transverse relaxation times are lower. This is visually illustrated by the corresponding difference maps below. This would indicate that the extra-axonal water *T*_2_ is lower than the axonal *T*_2_. However, this trend – although generally confirmed in most WM regions – is opposite to what found for the spherical mean *T*_2_ in the WM regions with high FA as illustrated by the corresponding difference maps of fig. 6. This inverted trend could be the effect of the presence of other compartments contributing to the low b-value spherical mean signal in addition to the isotropically-restricted one.

**Figure 6:**
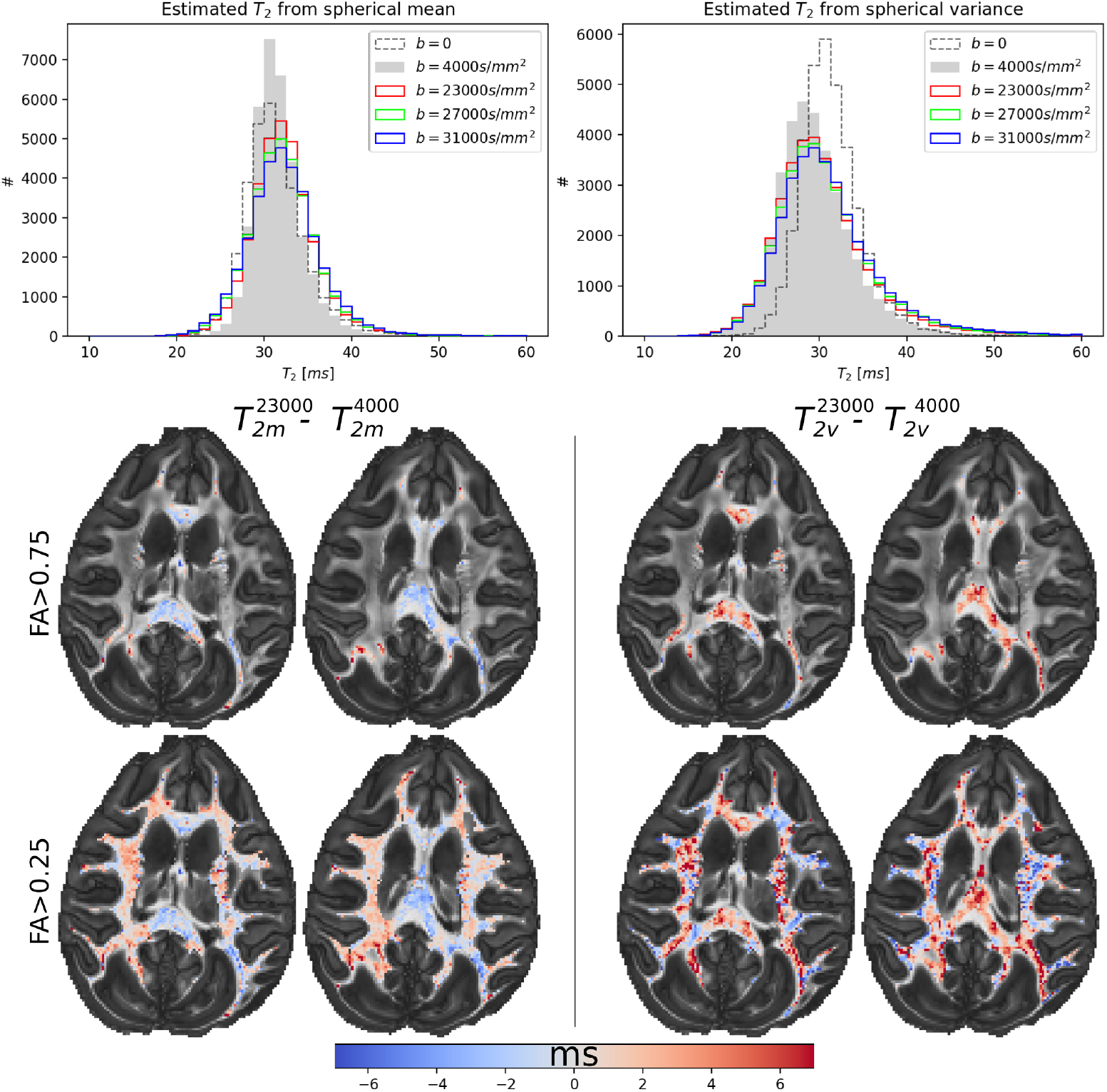
*Ex vivo* data. As the b-value increases, the least restricted anisotropic compartments (e.g. the extra-axonal one) cease to contribute to the estimated spherical variance *T*_2v_ (right). Changes between the gray histogram (step-filled) and the colored ones (step) are then only determined by the disappearance of the least restricted tissue components from the signal. A similar trend is observed for the spherical mean estimates (left) histograms which is however inverted in the high FA regions. The histograms for *b* > 0 were calculated only accounting for WM voxels where only one fiber bundle was detected.

### 3.3. *In vivo* human data

The analysis performed for the *ex vivo* monkey data was replicated for the *in vivo* human data. In particular, the derived *T*_2_ maps are shown in fig. 7, using a white matter mask defined over the voxels where FA > 0.25. We note a prevalence of voxels where the spherical variance *T*_2v_ (third column) is higher than both the average *b* = 0 *T*_2_ (first column) and the spherical mean *T*_2m_ (second column), as also illustrated by the difference maps in the fourth column of the figure. In some voxels, the extent of the differences approaches 50 ms (fourth column). By comparing the high b-value spherical variance *T*_2_ estimates with those based on the spherical mean it is possible to deduce that isotropically-restricted compartments have lower *T*_2_ values than the axonal compartments in the red-colored regions of the maps illustrated in the fourth column of fig. 7. This trend, however, shows local differences.

**Figure 7:**
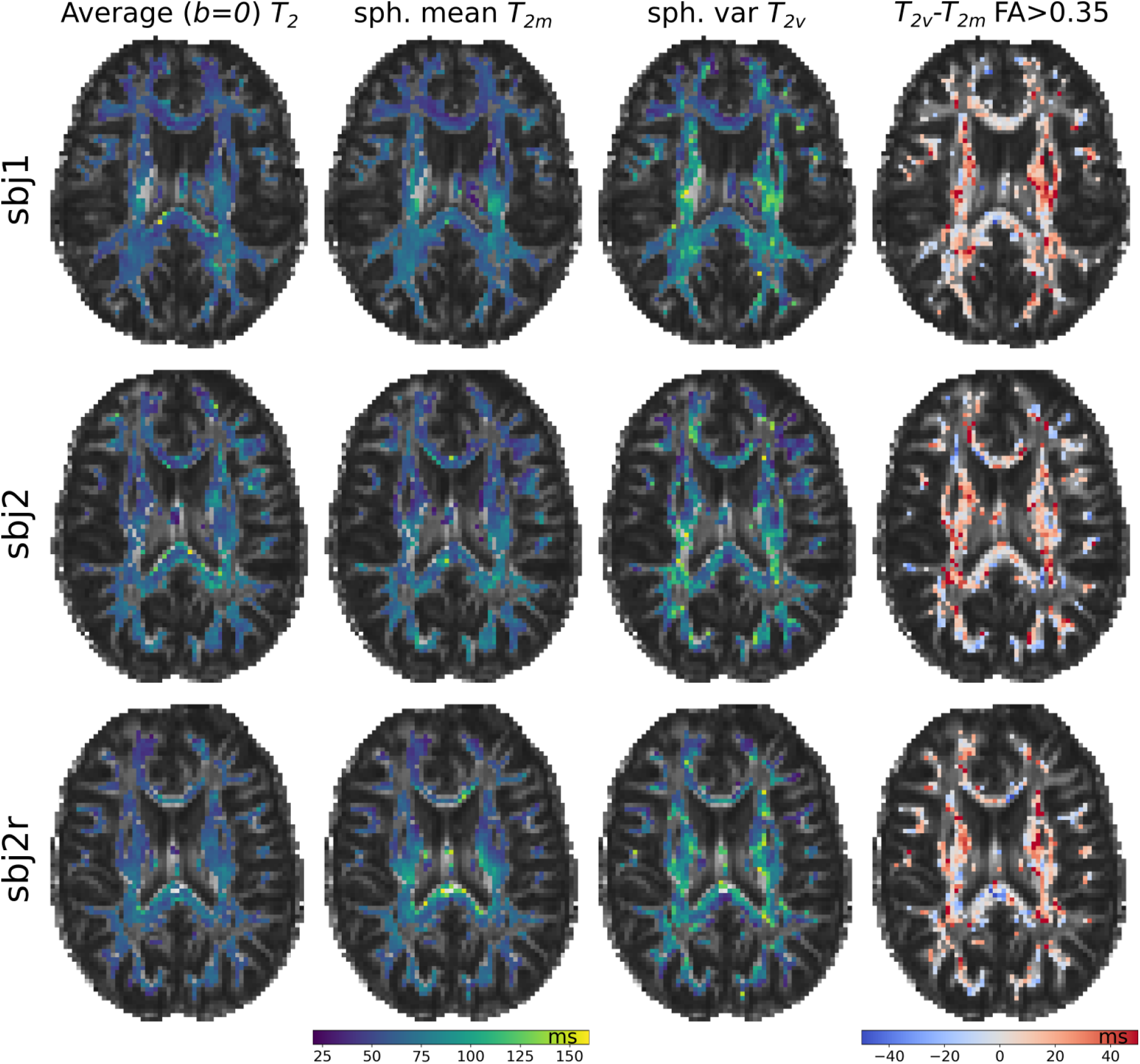
*In vivo* human data. Estimates of *T*_2_ for regions with FA > 0.35 for the different acquisitions. The smoothed versions of *T*_2v_ maps are reported. Rows correspond to different subjects and/or repetitions.

The *T*_2v_ estimates have been obtained from spherical variances calculated from the spherical harmonics coefficients for an expansion up to order *L* = 8 without applying any regularization. The spherical variance *T*_2_ estimates show a lower stability than those based on the spherical mean, which can be attributed to the concomitant effects of low (noise-free) spherical variance and to residual noise, as described previously. To cope with this, we have attempted a smoothing technique where for each voxel the two neighboring voxels with the most similar signal (according to the mean squared differences) were additionally used to fit the voxel’s *T*_2v_ value, similarly to what done in Alexander et al. (2010). Figures 8 and 10 illustrate the results of the quantitative analysis. We observe that the smoothing reduces the variability of the estimates although substantially preserving the shape of the distribution.

**Figure 8:**
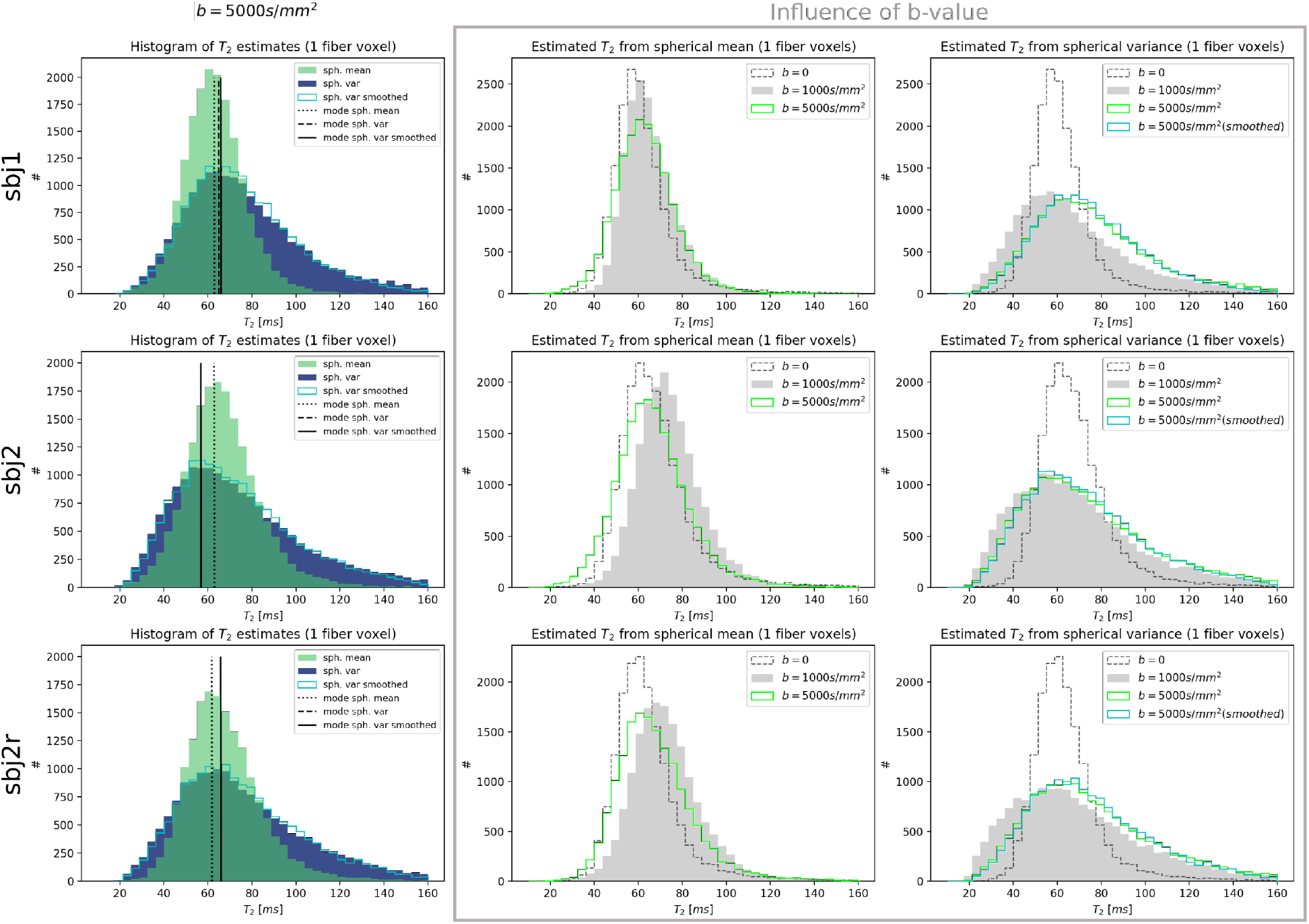
*In vivo* human data. Histograms of *T*_2_ estimates for the different acquisitions. On the right, histograms showing the influence of b-value on the estimates (spherical mean *T*_2m_ in the left column and spherical variance *T*_2v_ in the right one). Note that for subject 2 the mode of the spherical variance is hidden behind that of its smoothed version.

The right-side panel of fig. 8 shows the histograms illustrating the effect of increasing b-value for the spherical mean and spherical variance *T*_2_ estimates. These trends show similarities with the *ex vivo* results, as also illustrated for subject 1 in fig. 9, although the variability of the estimates does not allow us to draw more general conclusions (see supplemental fig. S2).

**Figure 9:**
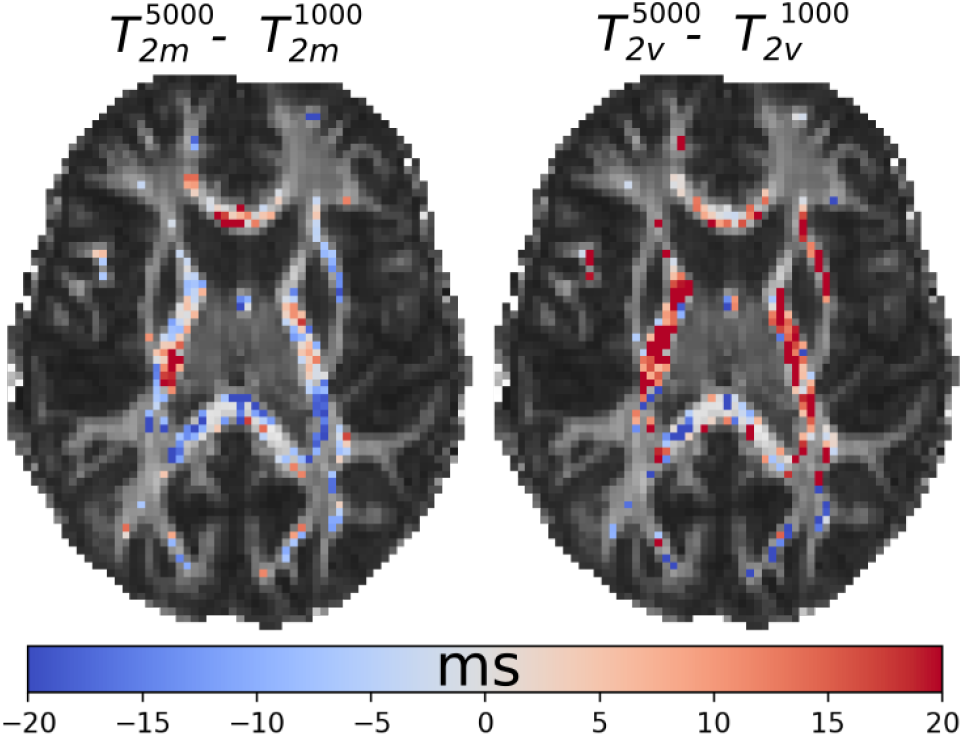
*In vivo* human data (sbj1). Differences between estimates calculated at *b* = 5000 s/mm^2^ and *b* = 1000 s/mm^2^ for the dataset corresponding to subject 1, in the case of spherical mean (left) and spherical variance (right) estimators. A more complete view is illustrated in the supplemental fig. S2.

The *T*_2_ estimates have a dependency on the anisotropy (see the influence of FA in fig. 10). Compared to the *ex vivo* case, where the spherical mean *T*_2_ estimates decrease with FA, for the *in vivo* case all the *T*_2_ estimates have an increasing trend with FA. As for the *ex vivo* case, the panel reporting the effect of the angle of the fiber with respect to the B0 field direction illustrates decreasing trends of the *T*_2_ estimates as the angle increases. It has to be noted that in fig. 10 the trends are indicated by the mode of the distributions, since the spherical variance distributions are asymmetric, as illustrated in fig. 8, which was not the case for the *ex vivo* results where the median of the distributions was a good estimate of the mode. The estimation of the mode is based on a cubic interpolation of the histogram to find the peak of the distributions: these estimates are less stable and precise than the median and this can be reflected in seemingly more unstable lines.

**Figure 10:**
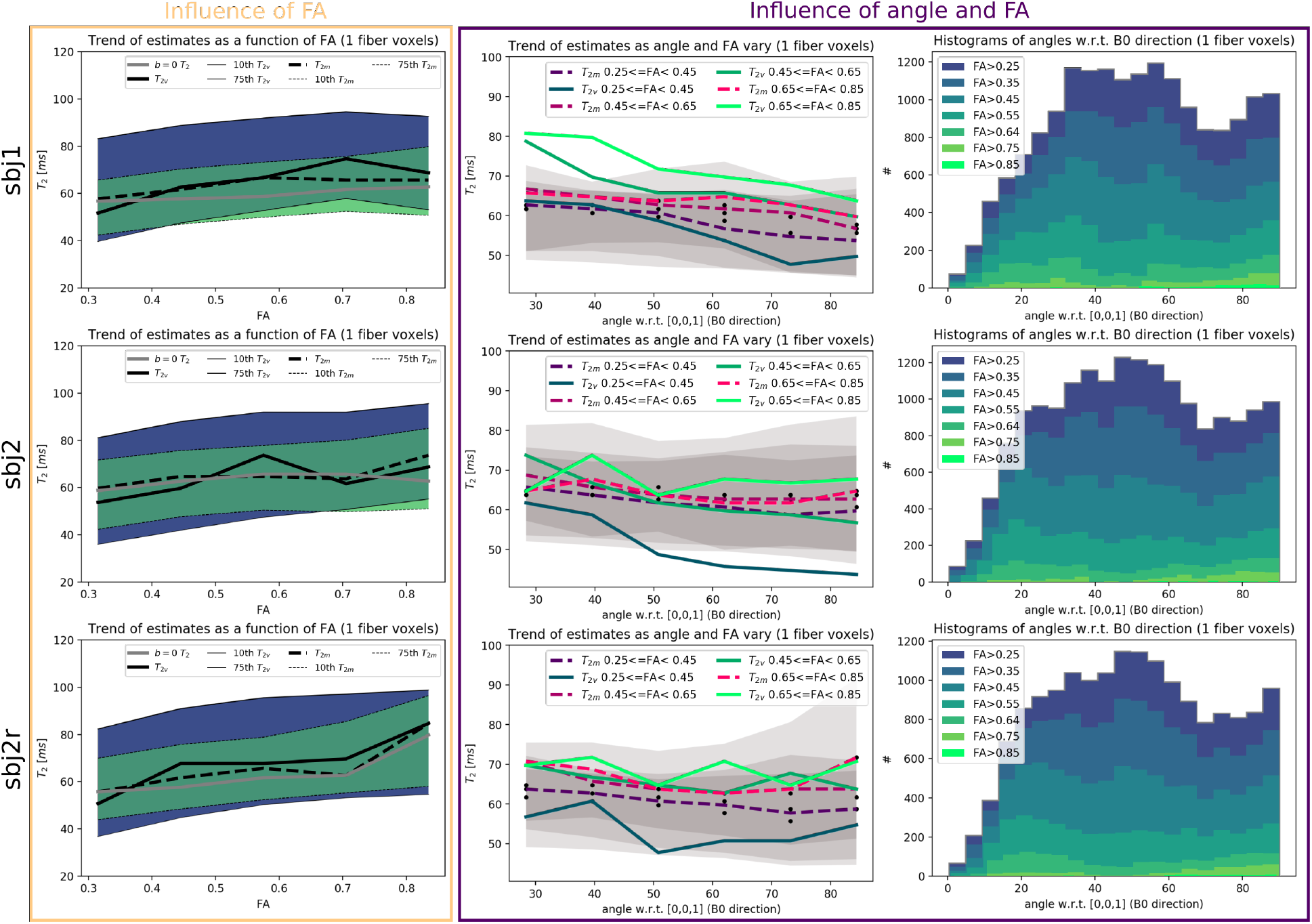
*In vivo* human data. Trend of the estimates as a function of FA on the left, and as a function of FA and of the detected angle (DTI) between the fibers and the B0 field direction on the right. In all plots, lines report the estimated mode of the distributions (except from those reporting the percentiles).

Although the histograms in fig. 8 show that the distributions of the different estimates of *T*_2_ are different, the panel about the influence of FA in fig. 10 illustrates that the mode of the distributions are well-aligned and close to each other. This is the case also for the average (*b* = 0) *T*_2_ estimates.

## 4. Discussion

In the presence of an isotropically-restricted compartment in the white matter, the axonal *T*_2_ estimator based on the spherical variance of the strongly diffusion-weighted MRI signal provides an unbiased estimation of the axonal transverse relaxation time, as opposed to the estimator based on the spherical mean. Indeed, the spherical variance *T*_2_ estimator is designed to be insensitive to tissue structures characterized by an isotropic contribution to the diffusion-weighted MRI signal, which has been demonstrated on simulated data. In the high b-value regime, therefore, differences between the *T*_2_ estimates obtained from the spherical variance and from the spherical mean of the signal have to be attributed uniquely to the presence of isotropically-restricted compartments. Since the spherical mean and spherical variance *T*_2_ estimators provide different estimates of *T*_2_ in both *ex vivo* fixed tissue and *in vivo* human brain, it is not possible to discard the presence of a detectable isotropically-restricted signal contribution such as that possibly associated to the cell nuclei or vacuoles identified by Andersson et al. (2020). These results complement existing knowledge (Dhital et al., 2018; Veraart et al., 2020; Tax et al., 2020) and add to the discussion on the presence of the so-called “dot” compartment which is, per-definition, an extreme case of an isotropically-restricted compartment where water appears to be immobile. Our analysis, however, does not allow for assessing the eventual dot-like nature of the isotropically-restricted compartment.

The differences between the spherical variance and spherical mean *T*_2_ estimates were more uniform for the *ex vivo* fixed data collected with the pre-clinical 7T MRI system which provided a higher SNR compared to the 3T clinical system. The use of smoothing on the human data revealed substantially the same distributions of the spherical variance *T*_2_ estimates as to when smoothing was not used, which indicates that the amount of noise may not be the main factor determining the shape of the distribution. However, some doubts remain with regards to whether the differences between the spherical variance and the spherical mean *T*_2_ estimates are episodic, e.g. related to the specific local configurations of the white matter tissue or to the presence of residual partial voluming. These doubts are also motivated by the similarities of the mode values of the estimates illustrated, for the *in vivo* data, in figs. 8 and 10.

The detectability of the eventual presence of an isotropically-restricted compartment by means of the difference between spherical mean and spherical variance *T*_2_ estimates is only possible whenever such compartment is characterized by a different *T*_2_ compared to the axonal compartment. In the high SNR *ex vivo* data, the spherical mean and variance *T*_2_ estimates differed by a few milliseconds as illustrated in fig. 3. If differences of similar extent were to be expected for the *in vivo* human tissue it may be possible that these would be below the sensitivity achievable with the collected data. Small detectable absolute differences do not imply that the differences between the axonal and the isotropically-restricted *T*_2_ values are also small. As illustrated by the simulation results of fig. 2, the absolute differences observed between the spherical mean and spherical variance *T*_2_ values are lower than the actual difference between the compartmental *T*_2_ values and are modulated by the compartmental volume fractions.

In summary, based on the collected data, the comparison between the spherical variance and spherical mean*T*_2_ estimates in the strongly diffusion-weighted regime did not rule out the presence of a detectable isotropically-restricted compartment in the human white matter tissue, while it strongly hinted to its presence in fixed *ex vivo* tissue. The proposed spherical variance *T*_2_ estimator should however be considered for the more general problem of estimating the *T*_2_ of anisotropic structures and could be integrated in the optimization procedures of multi-compartmental biophysical models.

### 4.1. Sensitivity to residual noise

In the presence of an isotropically-restricted compartment in the white matter, the axonal *T*_2_ estimator based on the spherical variance of the diffusion-weighted MRI signal provides an unbiased estimation as opposed to the estimator based on the spherical mean. The use of the spherical variance poses however more challenges in terms of sensitivity to noise since the noise variance adds directly to the ground-truth spherical variance. The spherical mean benefits instead from a higher effective signal-to-noise ratio since it is the result of an averaging procedure. Nevertheless, the use of a spherical harmonics expansion can be beneficial even when regularization is not applied. Indeed, the approximation of the directional signal on a shell can help remove the influence of the residual noise after denoising. However, such residual noise is likely to be correlated spatially and along the various diffusion gradient directions which can be less than ideal for the assumptions behind the linear approximation of the signal with spherical harmonics. Moreover, as for the spherical mean *T*_2_ estimator, the presence of a residual Rician bias could affect also the estimated *T*_2_ values from the spherical variance estimator.

The simulations have shown that the stability of the spherical variance *T*_2_ estimates reduces with the increasing b-value and number of fibers crossing within a voxel (fig. 2). More precisely, the stability is related to the amount of spherical variance of the noise-free directional signal, which in turn depends on the above mentioned factors and on axonal orientational dispersion. All these factors make of the spherical variance *T*_2_ a low sensitivity estimator, especially when considering the high b-value regime, although it can offer some advantages in terms of insensitivity to partial voluming effects (which could be especially useful at low diffusion weightings). The presence of residual noise has repercussions on the intra- and inter-subject variability of the spherical variance estimates, which will need further assessment by using more subjects and different acquisition protocols, filed strengths, and MRI scanners. It is likely, however, that the accuracy and stability of the estimates will improve with the development of more performing denoising methods.

### 4.2. Directional dependence of *T*_2_ estimates

For both *ex vivo* (7T) and *in vivo* (3T) datasets we found that the axonal *T*_2_, estimated with either the spherical mean or the spherical variance estimators at high b-value, was correlated with the angle between the direction of the WM fiber bundle and the direction of the B0 field. In particular, the estimated *T*_2_ decreased with the increasing angle with lower values identified for a 90° angle (Sati et al., 2012). A similar directional dependence has also been illustrated by McKinnon and Jensen (2019) for the spherical mean of the strongly diffusion-weighted signal. Using relaxometry, Birkl et al. (2021) detected an analogous dependence of the *T*_2_ values of both myelin and axonal/extra-axonal water. However, using a tilted RF coil system Tax, Kleban, Chamberland, Baraković, Rudrapatna and Jones (2021) attributed this dependence to the extra-axonal compartment instead of the axonal one. If that is the case, then it would be necessary to reconsider which is the surviving compartment in the high b-value regime, the axonal or the extra-axonal one. The identification of the axonal compartment as the “more” restricted one, and of the extra-axonal compartment as the “less” restricted one, is based on common assumptions in the field as it is supported by theoretical considerations on the geometry of the axonal space, such as its “stick-like” interpretation. In fact, although there are geometrical arguments for the axonal compartment being the most restricted, the determination of which compartment’ signal “survives” at strong diffusion weightings should take in consideration other factors, such as the intrinsic diffusivities of the two media. The use of axonal and extra-axonal terminology in this work should therefore be taken with these considerations in mind. Even if the extra-axonal space is less restricted than the axonal one, it is still possible that with the adopted b-values there still are non-negligible contributions from the former to the measured signal.

### 4.3. Regional dependence of *T*_2_ estimates

Despite showing different overall trends, results on *ex vivo* fixed tissue and on *in vivo* tissue support the existence of a correlation between the measured fractional anisotropy and the estimated *T*_2_ which is not linked to the presence of the correlation between *T*_2_ and the angle that fibers make with the B0 field direction. At the end of sec. 3.2.2 we examined the possibility that the changes in anisotropy, and the correlated changes in *T*_2_, might be more related to the presence of isotropically-restricted compartments rather than to the change in axonal dispersion. This consideration was supported by the observation that in *ex vivo* data the spherical mean *T*_2m_ seemed to vary more than the spherical variance *T*_2v_ as a function of FA. However, the *in vivo* results in fig. 10 support the opposite analysis. While the logical step would be trying to correlate the changes in FA with the amount of axonal dispersion, the estimation of the axonal dispersion would be biased unless accounting for the presence of isotropic compartments. The simultaneous characterization of isotropically-restricted compartments and orientation dispersion is challenging and would require further research. Therefore, this remains an open question for future work to which multi-compartmental modeling and/or higher SNR *in vivo* data could help find an answer.

### 4.4. Non-ideal human *in vivo* data

It was not possible to specify the pulse gradient strength, duration, and separation of the PGSE sequence used in the clinical 3T scanner, like in the case of McKinnon and Jensen (2019). The sequence provided by the vendor adapts the pulse duration and separation to the specified echo time, thus changing the diffusion-weighting although providing the same b-value (e.g. 5000 s/mm^2^). This led us to adopt a small echo time spacing to keep the diffusion weighting between the two different acquisitions as homogeneous as possible. Considering a minimum echo time of 80 ms, the adopted echo time spacing of 9 ms provides an attenuation of the signal between 12 to 16.5% when considering an axonal *T*_2_ of 70 and 50 ms respectively, as compared to a 23 to 30% with a double spacing of 18 ms which is more similar to that used by McKinnon and Jensen (2019). The chosen echo time spacing, on the other hand, provided a higher effective SNR. Future work will need to address the PGSE parameters to obtain a homogeneous diffusion weighting across echo times, as done for the pre-clinical acquisition.

### 4.5. The assumption of a single *T*_2_ compartment

Reference has often been made to the estimation of a unique *T*_2_ value from signal containing contributions from multiple compartments, such as the case of the average (*b* = 0) *T*_2_. This was motivated by the observation in synthetic data that the presence of an additional compartment will shift the estimated *T*_2_ towards higher or lower values when having a higher or lower *T*_2_ compared to a reference compartment. It is however important to note that in these cases the estimated *T*_2_ should be considered as biased. This is indeed the reason for using the spherical variance estimator instead of that based on the spherical mean. Nevertheless, differences between *T*_2_ estimators (eventually biased) reveal that there are different compartmental contributions to the signal, which is the main method we used to assert the differences in the results. However, the meta-analysis about the relative order between the *T*_2_ values of the axonal, extra-axonal, and isotropically-restricted compartments should be considered as only indicative, and a more thorough assessment remains to be done perhaps with the use of multi-compartmental biophysical modeling across all the b-values (Veraart, Novikov and Fieremans, 2018). Moreover, the regional dependence of the differences between the average (*b* = 0) *T*_2_, the spherical mean *T*_2m_, and the *T*_2v_ makes it difficult to derive a general rule for the order of the compartmental transverse relaxation times.

### 4.6. The unknown impact of fixation

The microstructural properties of the *ex vivo* tissue could differ from those of the living tissue both due to the effects of the *post mortem* state and of the fixation. Andersson et al. (2020) suggest, for instance, that the fixation could be the cause of the vacuolation of the tissue. This could exacerbate the isotropic contributions to the diffusion-weighted signal compared to the natural contributions in the living tissue. However, the presence of cellular components in the living white matter tissue is well established. Moreover, the presence of cells and microglia can also be related to the occurrence of inflammation or other types of pathological conditions (Mammana, Fagone, Cavalli, Basile, Petralia, Nicoletti, Bramanti and Mazzon, 2018; He, Aznar, Siebner and Dyrby, 2021). Nevertheless, the results for fixed tissue cannot be directly compared to those *in vivo*. Indeed, Birkl et al. (2016) report an approximately 30% decrease of the *T*_2_ values in white matter due to the fixation involving the use of formalin. Use of saline washing of the tissue after fixation may however help to partly regain the *T*_2_ values in white matter (Leprince, Schmitt, Chaillou, Destrieux, Barantin, Vignaud, Rivière and Poupon, 2015; Dyrby, Innocenti, Bech and Lundell, 2018). Additionally, the effects of fixation may be inhomogeneous across tissue constituents. Fixation and *post mortem* may then explain the discrepancies between *ex vivo* and *in vivo* results, such as the decreasing and increasing trends of the *T*_2_ estimates as a function of FA.

### 4.7. Other diffusion-related considerations

The signal model assumed in eq. 3 offers a rather simplistic view of the diffusion process in the strongly diffusion-weighted regime. First of all, it assumes that the anisotropic signal contributions are uniquely determined by the axonal compartment. However, it is natural to expect that cell nuclei and vacuoles deviate from a perfect spherical shape, as also measured by Andersson et al. (2020). Nevertheless, the anisotropic contribution to the signal from these structures is expected to be negligible compared to that of the axons. Secondly, the model in eq. 3 assumes a tissue microstructure where barriers, such as cellular and axonal membranes, are completely impermeable to the passage of spin-bearing water particles. However, permeability is a relevant phenomenon (Sønderby, Lundell, Søgaard and Dyrby, 2014) especially when considering relatively long diffusion times (often associated to long echo times) such as those used for the *in vivo* clinical data (Li, Jiang, Xie, McIntyre, Gore and Xu, 2016). This is only the most evident reason why *T*_2_ estimates might be different with different acquisition setups where any of the pulse gradient strength, duration, separation, echo time, or repetition time should change. Interesting future work could therefore investigate the changes in the spherical mean and spherical variance *T*_2_ as a function of the different acquisition setups.

## 5. Conclusion

The calculation of *T*_2_ using the spherical variance of the diffusion-weighted MRI signal acquired over shells having different echo times provides sensitivity to only the anisotropic components of the tissue microstructure. When using strong diffusion weightings that suppress the signal coming from the less restricted diffusion compartments of the microstructure within a voxel, such as the extra-axonal space, the spherical variance *T*_2_ estimates are only influenced by the axonal (or the more restricted anisotropic) compartment while the spherical mean *T*_2_ estimates would still contain contributions from isotropically-restricted compartments. The results of the comparison between spherical variance and spherical mean *T*_2_ estimates do not allow us to discard the presence of one or more MRI-visible isotropically-restricted signal compartments in the white matter. A high signal-to-noise ratio of the acquired signal is crucial for using the spherical variance to estimate the axonal *T*_2_, and the improvement of denoising methods will allow for more robust estimates in the future. The use of spherical variance *T*_2_ estimates could be a complementary yet fundamental asset in the characterization of the white matter tissue and for microstructural modeling, with foreseeable applications in the detection and characterization of pathology.

## Acknowledgments

We thank Carmen Moreno Genis for facilitating the acquisition of the clinical MRI images.

## A. Spherical harmonics representation

The adopted spherical harmonics representation is

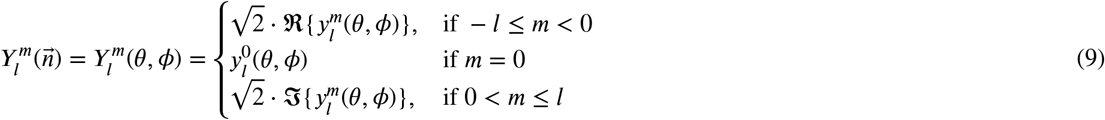

with *θ* ∈ [0, *π*] and *ϕ* ∈ [0, 2*π*] being the spherical coordinates for the vector 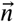, where the symbols ℜ and 𝔍 indicate the real and imaginary parts respectively, and where

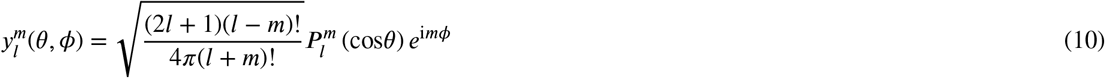

with 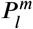 being the associated Legendre polynomials and i the imaginary unit.

## B. Naive estimators

We compare two different naive estimators, one based on the average of the *T*_2_ estimates across all the diffusion gradient directions and one based instead on the median. Let us define the signal

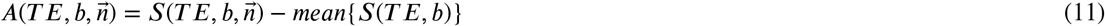

then

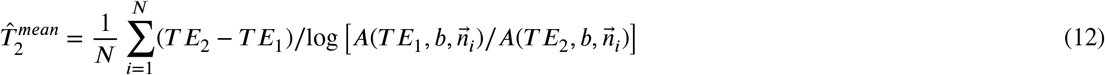

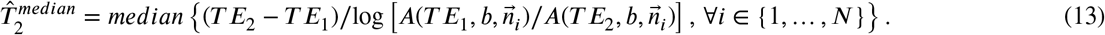

The maps obtained with the different estimators are reported in fig. 11, while the corresponding histograms are shown in fig. 12. For the directional median estimator, the directions corresponding to a measured *T*_2_ decay outside the [10,100] ms range were excluded from the final *T*_2_ estimates to improve stability. Similarly, for the directional mean estimator different ranges were tested: [10, 70],[10, 100], and [10, 200] ms.

**Figure 11:**
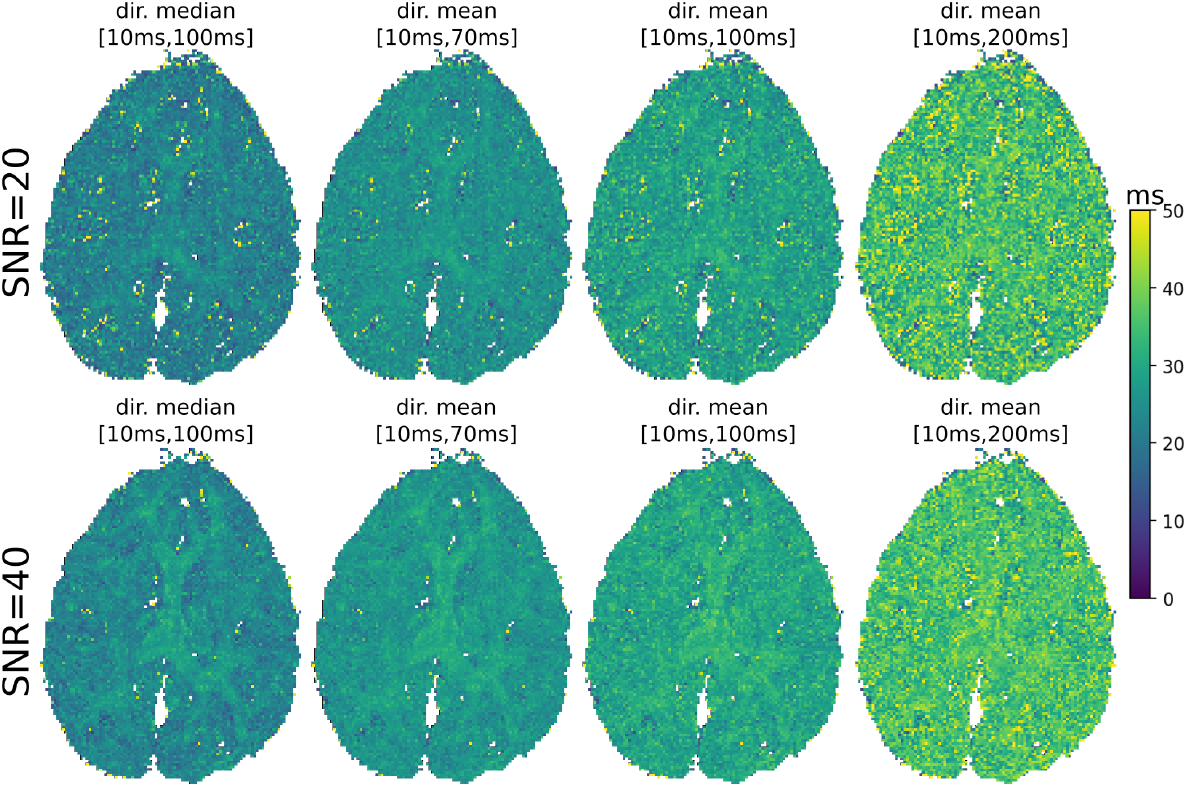
Synthetic data. Maps of axonal *T*_2_ estimates (ms) calculated at *b* = 23000 s/mm^2^ for SNR 20 and 40 on simulated data with the naive directional median and mean estimators. For all, the range of thresholded *T*_2_ values is specified in square brackets. The ground-truth axonal *T*_2_ was 30 ms.

**Figure 12:**
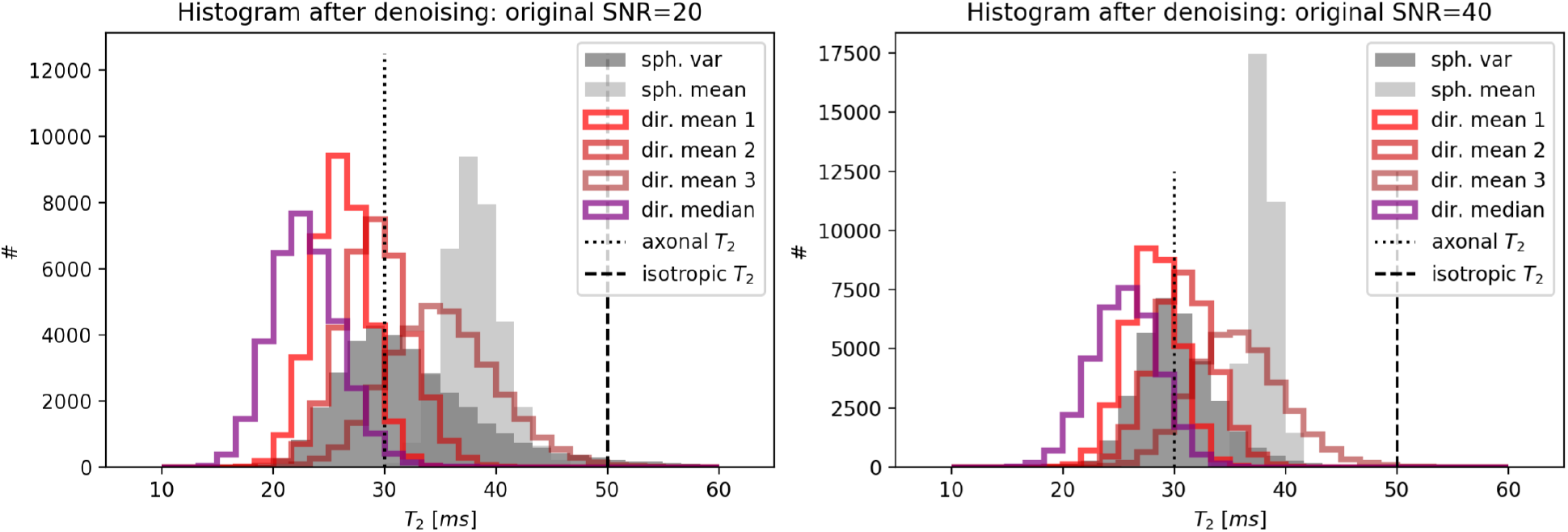
Synthetic data. Performance of the naive estimators compared to the spherical variance and spherical mean ones. The numbers 1, 2, and 3 for the directional mean estimates correspond to different admissible ranges for the *T*_2_ values similarly to what indicated in the corresponding maps of fig. 11. In particular 1=[10, 70 ms], 2=[10, 100 ms], and 3=[10, 200 ms]. Similarly, for the directional median estimator the range was fixed to [10, 100 ms].

The results indicated that while the directional median estimator shows better stability than that based on the spherical variance, it also shows lower accuracy leading to slightly more biased estimates. The directional mean estimator was instead found to be very sensitive to the imposed interval bounds. These results suggest that in order to estimate the axonal transverse relaxation time it is better to use the spherical variance estimator.

## Footnotes

1 https://www.mrtrix.org/

2 https://numpy.org/

3 https://www.scipy.org/

4 https://matplotlib.org/

5 https://github.com/AthenaEPI/dmipy

6 https://dipy.org/

## References

Alexander, D.C., Dyrby, T.B., Nilsson, M., Zhang, H., 2019. Imaging brain microstructure with diffusion MRI: practicality and applications. NMR in Biomedicine 32, e3841.

Alexander, D.C., Hubbard, P.L., Hall, M.G., Moore, E.A., Ptito, M., Parker, G.J., Dyrby, T.B., 2010. Orientationally invariant indices of axon diameter and density from diffusion MRI. Neuroimage 52, 1374–1389.

Andersson, J.L., Skare, S., Ashburner, J., 2003. How to correct susceptibility distortions in spin-echo echo-planar images: application to diffusion tensor imaging. Neuroimage 20, 870–888.

Andersson, J.L., Sotiropoulos, S.N., 2016. An integrated approach to correction for off-resonance effects and subject movement in diffusion mr imaging. Neuroimage 125, 1063–1078.

Andersson, M., Kjer, H.M., Rafael-Patino, J., Pacureanu, A., Pakkenberg, B., Thiran, J.P., Ptito, M., Bech, M., Dahl, A.B., Dahl, V.A., et al., 2020. Axon morphology is modulated by the local environment and impacts the noninvasive investigation of its structure–function relationship. Proceedings of the National Academy of Sciences 117, 33649–33659.

Barakovic, M., Tax, C.M., Rudrapatna, U., Chamberland, M., Rafael-Patino, J., Granziera, C., Thiran, J.P., Daducci, A., Canales-Rodríguez, E.J., Jones, D.K., 2021. Resolving bundle-specific intra-axonal T2 values within a voxel using diffusion-relaxation tract-based estimation. NeuroImage 227, 117617.

Basser, P.J., Mattiello, J., LeBihan, D., 1994. Estimation of the effective self-diffusion tensor from the nmr spin echo. Journal of Magnetic Resonance, Series B 103, 247–254.

Birkl, C., Doucette, J., Fan, M., Hernández-Torres, E., Rauscher, A., 2021. Myelin water imaging depends on white matter fiber orientation in the human brain. Magnetic resonance in medicine 85, 2221–2231.

Birkl, C., Langkammer, C., Golob-Schwarzl, N., Leoni, M., Haybaeck, J., Goessler, W., Fazekas, F., Ropele, S., 2016. Effects of formalin fixation and temperature on mr relaxation times in the human brain. NMR in Biomedicine 29, 458–465.

Descoteaux, M., Angelino, E., Fitzgibbons, S., Deriche, R., 2007. Regularized, fast, and robust analytical q-ball imaging. Magnetic resonance in medicine 58, 497–510.

Dhital, B., Kellner, E., Kiselev, V.G., Reisert, M., 2018. The absence of restricted water pool in brain white matter. Neuroimage 182, 398–406.

Dyrby, T.B., Baaré, W.F., Alexander, D.C., Jelsing, J., Garde, E., Søgaard, L.V., 2011. An ex vivo imaging pipeline for producing high-quality and high-resolution diffusion-weighted imaging datasets. Human brain mapping 32, 544–563.

Dyrby, T.B., Innocenti, G.M., Bech, M., Lundell, H., 2018. Validation strategies for the interpretation of microstructure imaging using diffusion MRI. Neuroimage 182, 62–79.

Ferizi, U., Schneider, T., Panagiotaki, E., Nedjati-Gilani, G., Zhang, H., Wheeler-Kingshott, C.A., Alexander, D.C., 2014. A ranking of diffusion MRI compartment models with in vivo human brain data. Magnetic resonance in medicine 72, 1785–1792.

Fick, R.H., Wassermann, D., Deriche, R., 2019. The dmipy toolbox: Diffusion MRI multi-compartment modeling and microstructure recovery made easy. Frontiers in neuroinformatics 13, 64.

Foi, A., 2011. Noise estimation and removal in mr imaging: The variance-stabilization approach, in: 2011 IEEE International symposium on biomedical imaging: from nano to macro, IEEE. pp. 1809–1814.

Garyfallidis, E., Brett, M., Amirbekian, B., Rokem, A., Van Der Walt, S., Descoteaux, M., Nimmo-Smith, I., 2014. Dipy, a library for the analysis of diffusion MRI data. Frontiers in neuroinformatics 8, 8.

Gavish, M., Donoho, D.L., 2017. Optimal shrinkage of singular values. IEEE Transactions on Information Theory 63, 2137–2152.

Harris, C.R., Millman, K.J., van der Walt, S.J., Gommers, R., Virtanen, P., Cournapeau, D., Wieser, E., Taylor, J., Berg, S., Smith, N.J., Kern, R., Picus, M., Hoyer, S., van Kerkwijk, M.H., Brett, M., Haldane, A., Fernández del Río, J., Wiebe, M., Peterson, P., Gérard-Marchant, P., Sheppard, K., Reddy, T., Weckesser, W., Abbasi, H., Gohlke, C., Oliphant, T.E., 2020. Array programming with NumPy. Nature 585, 357–362.

He, Y., Aznar, S., Siebner, H.R., Dyrby, T.B., 2021. In vivo tensor-valued diffusion MRI of focal demyelination in white and deep grey matter of rodents. NeuroImage: Clinical 30, 102675.

Hunter, J.D., 2007. Matplotlib: A 2d graphics environment. Computing in Science & Engineering 9, 90–95.

Jelescu, I.O., Veraart, J., Fieremans, E., Novikov, D.S., 2016. Degeneracy in model parameter estimation for multi-compartmental diffusion in neuronal tissue. NMR in Biomedicine 29, 33–47.

Jensen, J.H., Glenn, G.R., Helpern, J.A., 2016. Fiber ball imaging. Neuroimage 124, 824–833.

Jeurissen, B., Tournier, J.D., Dhollander, T., Connelly, A., Sijbers, J., 2014. Multi-tissue constrained spherical deconvolution for improved analysis of multi-shell diffusion MRI data. NeuroImage 103, 411–426.

Jones, D.K., Horsfield, M.A., Simmons, A., 1999. Optimal strategies for measuring diffusion in anisotropic systems by magnetic resonance imaging. Magnetic Resonance in Medicine: An Official Journal of the International Society for Magnetic Resonance in Medicine 42, 515–525.

Kellner, E., Dhital, B., Kiselev, V.G., Reisert, M., 2016. Gibbs-ringing artifact removal based on local subvoxel-shifts. Magnetic resonance in medicine 76, 1574–1581.

Leprince, Y., Schmitt, B., Chaillou, É., Destrieux, C., Barantin, L., Vignaud, A., Rivière, D., Poupon, C., 2015. Optimization of sample preparation for MRI of formaldehyde-fixed brains, in: Proceedings of the 23rd Annual Meeting of ISMRM, Toronto.

Li, H., Jiang, X., Xie, J., McIntyre, J.O., Gore, J.C., Xu, J., 2016. Time-dependent influence of cell membrane permeability on mr diffusion measurements. Magnetic resonance in medicine 75, 1927–1934.

Ma, X., Uğurbil, K., Wu, X., 2020. Denoise magnitude diffusion magnetic resonance images via variance-stabilizing transformation and optimal singular-value manipulation. Neuroimage 215, 116852.

Mammana, S., Fagone, P., Cavalli, E., Basile, M.S., Petralia, M.C., Nicoletti, F., Bramanti, P., Mazzon, E., 2018. The role of macrophages in neuroinflammatory and neurodegenerative pathways of alzheimer’s disease, amyotrophic lateral sclerosis, and multiple sclerosis: Pathogenetic cellular effectors and potential therapeutic targets. International journal of molecular sciences 19, 831.

McKinnon, E.T., Jensen, J.H., 2019. Measuring intra-axonal T2 in white matter with direction-averaged diffusion MRI. Magnetic resonance in medicine 81, 2985–2994.

Panagiotaki, E., Schneider, T., Siow, B., Hall, M.G., Lythgoe, M.F., Alexander, D.C., 2012. Compartment models of the diffusion mr signal in brain white matter: a taxonomy and comparison. Neuroimage 59, 2241–2254.

Ramanna, S., Moss, H.G., McKinnon, E.T., Yacoub, E., Helpern, J.A., Jensen, J.H., 2020. Triple diffusion encoding MRI predicts intra-axonal and extra-axonal diffusion tensors in white matter. Magnetic resonance in medicine 83, 2209–2220.

Sati, P., Silva, A.C., van Gelderen, P., Gaitan, M.I., Wohler, J.E., Jacobson, S., Duyn, J.H., Reich, D.S., 2012. In vivo quantification of T2^*^ anisotropy in white matter fibers in marmoset monkeys. Neuroimage 59, 979–985.

Smith, S.M., Jenkinson, M., Woolrich, M.W., Beckmann, C.F., Behrens, T.E., Johansen-Berg, H., Bannister, P.R., De Luca, M., Drobnjak, I., Flitney, D.E., et al., 2004. Advances in functional and structural mr image analysis and implementation as fsl. Neuroimage 23, S208–S219.

Sønderby, C.K., Lundell, H.M., Søgaard, L.V., Dyrby, T.B., 2014. Apparent exchange rate imaging in anisotropic systems. Magnetic resonance in medicine 72, 756–762.

Stejskal, E.O., Tanner, J.E., 1965. Spin diffusion measurements: spin echoes in the presence of a time-dependent field gradient. The journal of chemical physics 42, 288–292.

Storn, R., Price, K., 1997. Differential evolution–a simple and efficient heuristic for global optimization over continuous spaces. Journal of global optimization 11, 341–359.

Tax, C.M., Szczepankiewicz, F., Nilsson, M., Jones, D.K., 2020. The dot-compartment revealed? diffusion MRI with ultra-strong gradients and spherical tensor encoding in the living human brain. NeuroImage, 116534.

Tax, C.M.W., Kleban, E., Chamberland, M., Baraković, M., Rudrapatna, U., Jones, D.K., 2021. Measuring compartmental T2-orientational dependence in human brain white matter using a tiltable RF coil and diffusion-T2 correlation MRI. NeuroImage 236, 117967.

Tournier, J.D., Calamante, F., Connelly, A., 2007. Robust determination of the fibre orientation distribution in diffusion MRI: non-negativity constrained super-resolved spherical deconvolution. Neuroimage 35, 1459–1472.

Veraart, J., Novikov, D.S., Fieremans, E., 2018. Te dependent diffusion imaging (teddi) distinguishes between compartmental T2 relaxation times. Neuroimage 182, 360–369.

Veraart, J., Nunes, D., Rudrapatna, U., Fieremans, E., Jones, D.K., Novikov, D.S., Shemesh, N., 2020. Noninvasive quantification of axon radii using diffusion MRI. Elife 9, e49855.

Virtanen, P., Gommers, R., Oliphant, T.E., Haberland, M., Reddy, T., Cournapeau, D., Burovski, E., Peterson, P., Weckesser, W., Bright, J., van der Walt, S.J., Brett, M., Wilson, J., Millman, K.J., Mayorov, N., Nelson, A.R.J., Jones, E., Kern, R., Larson, E., Carey, C.J., Polat, i., Feng, Y., Moore, E.W., VanderPlas, J., Laxalde, D., Perktold, J., Cimrman, R., Henriksen, I., Quintero, E.A., Harris, C.R., Archibald, A.M., Ribeiro, A.H., Pedregosa, F., van Mulbregt, P., SciPy 1.0 Contributors, 2020. SciPy 1.0: Fundamental Algorithms for Scientific Computing in Python. Nature Methods 17, 261–272.

White, N.S., McDonald, C.R., Farid, N., Kuperman, J., Karow, D., Schenker-Ahmed, N.M., Bartsch, H., Rakow-Penner, R., Holland, D., Shabaik, A., et al., 2014. Diffusion-weighted imaging in cancer: physical foundations and applications of restriction spectrum imaging. Cancer research 74, 4638–4652.

Zeng, Q., Shi, F., Zhang, J., Ling, C., Dong, F., Jiang, B., 2018. A modified tri-exponential model for multi-b-value diffusion-weighted imaging: a method to detect the strictly diffusion-limited compartment in brain. Frontiers in neuroscience 12, 102.

Zucchelli, M., Deslauriers-Gauthier, S., Deriche, R., 2020. A computational framework for generating rotation invariant features and its application in diffusion MRI. Medical image analysis 60, 101597.

