## Supplementary figures S1 and S2 for "Axonal *T*_2_ estimation using the spherical variance of the strongly diffusion-weighted MRI signal"

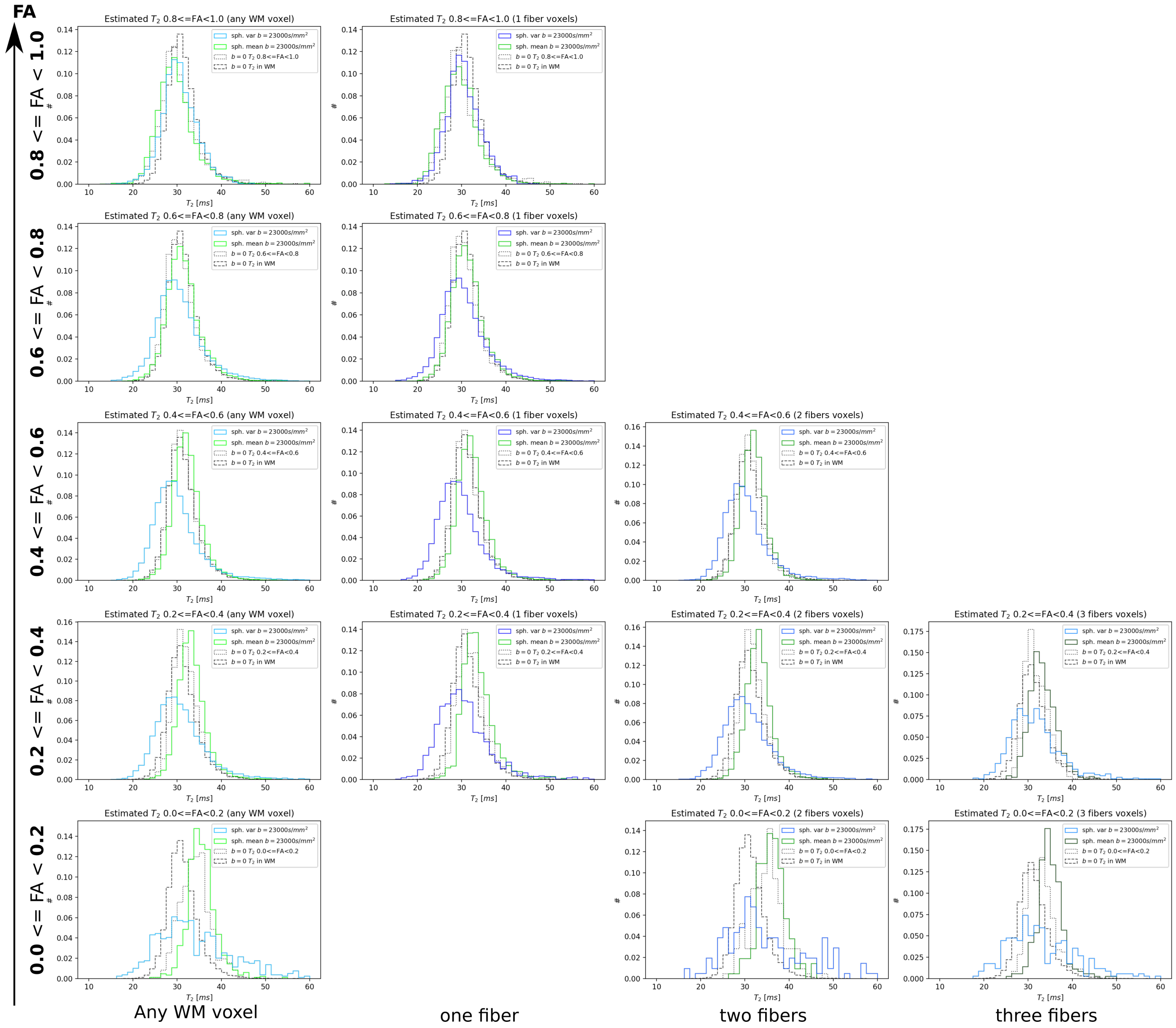


S1. *Ex vivo* data. Density histograms of axonal $T_2$ estimates for different ranges of FA (increasing along the vertical dimension) when considering all the white matter voxels (left column) or voxels with increasing number of detected fiber directions (horizontal dimension from left to right). ``Missing" entries are due to the insufficient number of voxels fulfilling the criteria, e.g. the simultaneous presence of three fibers and high FA values.

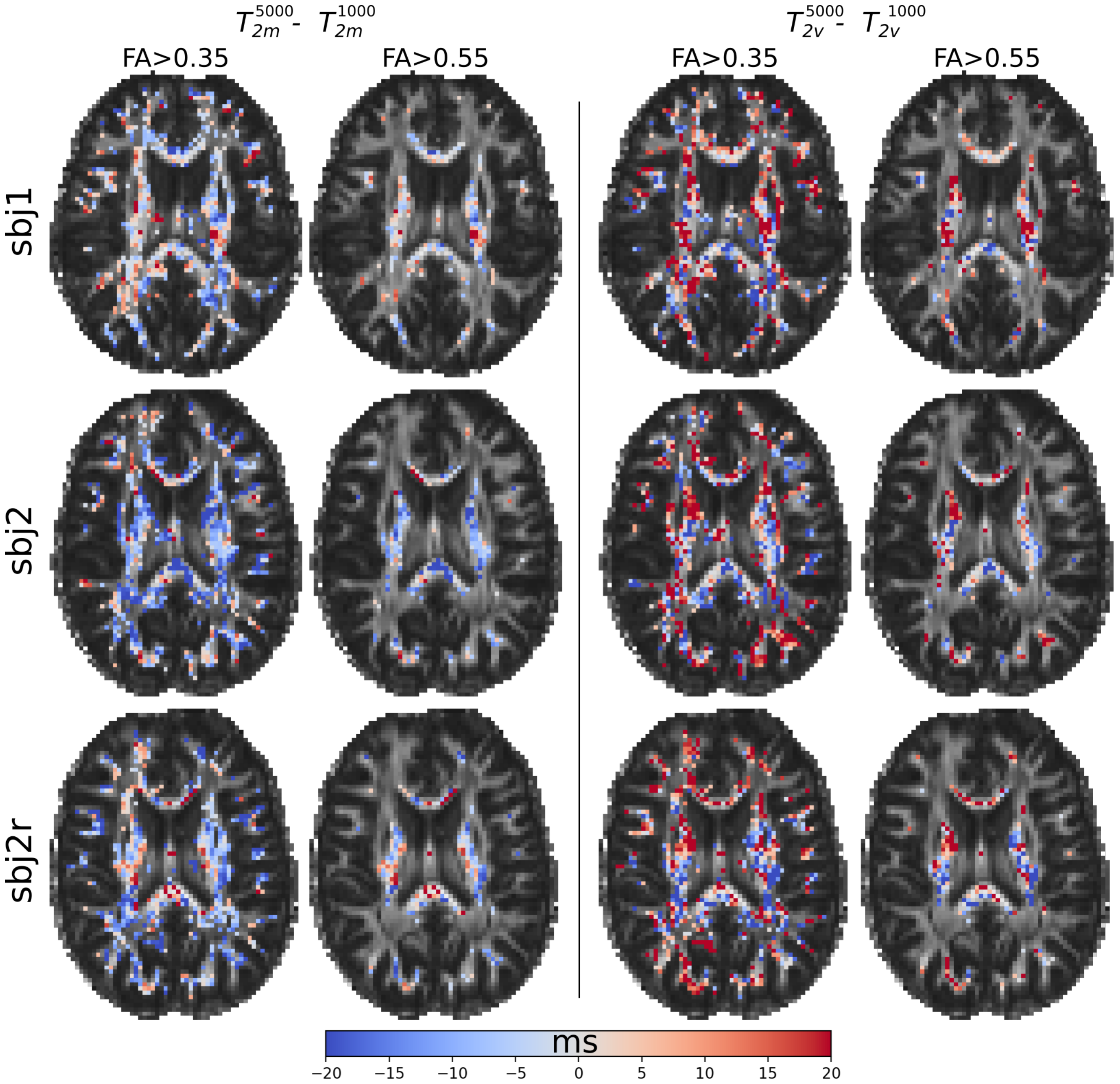


S2. *In vivo* data. Differences between estimates calculated at b=5000 s/mm$^2$ and b=1000 s/mm$^2$ for the various datasets, in the case of spherical mean (left) and spherical variance (right) estimators.
